# Molecular basis for the acid initiated uncoating of human enterovirus D68

**DOI:** 10.1101/361030

**Authors:** Yue Liu, Ju Sheng, Michael G. Rossmann

**Affiliations:** Department of Biological Sciences, Hockmeyer Hall of Structural Biology, 240 South Martin Jischke Drive, Purdue University, West Lafayette, IN 47907, USA

**Keywords:** Enterovirus D68, cryo-electron microscopy, virus uncoating, conformational changes, acidification

## Abstract

Enterovirus D68 (EV-D68) belongs to a group of enteroviruses that contain a single positive-sense RNA genome surrounded by an icosahedral capsid. Like common cold viruses, EV-D68 mainly causes respiratory infections and is acid labile. The molecular mechanism by which the acid sensitive EV-D68 virions uncoat and deliver their genome into a host cell is unknown. Using cryo-electron microscopy (cryo-EM), we have determined the structures of the full native virion and an uncoating intermediate (the A(altered)-particle) of EV-D68 at 2.2 Å and 2.7 Å resolution. These structures showed that acid treatment of EV-D68 leads to particle expansion, externalization of the viral protein VP1 N-termini from the capsid interior, and formation of pores around the icosahedral two-fold axes through which the viral RNA can exit. Moreover, because of the low stability of EV-D68 at neutral pH, cryo-EM analyses of a mixed population of particles demonstrated the involvement of multiple structural intermediates during virus uncoating. Among these, a previously undescribed state, the expanded (“E1”) particle, shows a majority of internal regions (e.g, the VP1 N-termini) to be ordered as in the full native virion. Thus, the E1 particle acts as an intermediate in the transition from full native virions to A-particles. Molecular determinants, including a histidine-histidine pair near the two-fold axes, were identified that facilitate this transition under acidic conditions. Thus, the present work delineates the pathway of EV-D68 uncoating and provides the molecular basis for the acid lability of EV-D68 and of the related common cold viruses.

**Significance Statement:** Enterovirus D68 (EV-D68) is an emerging pathogen that primarily causes childhood respiratory infections and is linked to neurological diseases. It was unclear how the virus uncoats and delivers its genome into a host cell to establish viral replication. Using high resolution cryo-electron microscopy, we showed that acid induces structural rearrangements of EV-D68 to initiate genome release from the virus. Structural analyses delineated a viral uncoating pathway that involves multiple distinct conformational states. Particularly, the structure of a previously unknown uncoating intermediate enabled the identification of molecular determinants that facilitate EV-D68 uncoating in an acidic environment. These results advance the knowledge of cell entry of EV-D68 and open up possibilities for developing antiviral therapeutics that impede structural rearrangements of the virus.

## Introduction

Enteroviruses (EVs) are a genus of single-stranded RNA viruses with a positive sense RNA genome surrounded by an icosahedral capsid shell (1). EVs from seven species, EV-A, EV-B, EV-C, EV-D, rhinovirus(RV)-A, RV-B and RV-C, are causative agents of a variety of human diseases (2, 3). These viruses include polioviruses (PVs), coxsackieviruses (CVs), RVs, EV-A71, and EV-D68. Among these, EV-D68 is a globally emerging human pathogen that mainly causes respiratory infections in young children (4-7). It has also been closely linked to neurological diseases (7-10). The development of effective vaccines and antiviral treatments against EV-D68 has been difficult due to limited knowledge of the molecular mechanisms of virus infection. In particular, despite recent progress in studying receptor-dependent cell entry of EV-D68 (11-16), it remains unclear how the virus uncoats and delivers its genome into host cells.

Enterovirus capsids are assembled from 60 copies of viral proteins VP1, VP2, VP3, and VP4 with pseudo T=3 icosahedral symmetry (17, 18). The VP1, VP2, and VP3 subunits, each having an eight-stranded β-barrel “jelly roll” fold, form the icosahedral shell with an outer diameter of ~300 Å (17, 18). The capsid inner surface is decorated by 60 copies of VP4, together with the N-termini of VP1, VP2, and VP3. During EV infection, host factors, such as cellular receptors and endosomal acidification, trigger EV uncoating by altering the capsid structure (19, 20). The uncoating process has been proposed to proceed via a structural intermediate, the A(altered)-particle (21-23), characterized by loss of VP4 and by externalization of the VP1 N-terminal residues (21, 24-27). These structural changes precede viral penetration of the membranes of intracellular compartments (21, 28, 29). This facilitates genome release from the A-particles into the cytosol of host cells and the production of emptied particles (22, 30, 31).

EV-D68 shares features, including acid lability, with rhinoviruses that are respiratory viruses from the species RV-A, RV-B, and RV-C (4, 12). However, sequence comparisons have shown that EV-D68 is more closely related to members of the species EV-A, EV-B, and EV-C (4). These viruses, as exemplified by polioviruses, are resistant to the acidic environment of the human gastrointestinal tract (1). It has been established that acid treatment of RVs *in vitro* causes structural alterations of the virus and often leads to the formation of uncoating intermediates, including A-particles and emptied particles (32-34). Moreover, endosomal acidification acts as an important cue for RV uncoating in host cells (19, 22, 35). By analogy with RVs (36), it is probable that acid triggers EV-D68 uncoating.

Here we report that an EV-D68 isolate from the 2014 outbreak in the United States is particularly sensitive to acid. Acid treatment causes dramatic conformational changes of full native virions to form A-particles. Moreover, cryo-electron microscopic (cryo-EM) analyses of the virus at neutral pH show that EV-D68 uncoating proceeds via multiple structural intermediates. These include a previously unknown structural state, the expanded 1 (E1) particle, which retains ordered VP1 N-terminal residues and ordered VP4. Structural determinants, particularly a histidine-histidine pair, have been identified that promote irreversible conversion of the virus to the A-particle state via the E1 particle intermediate. These observations provide a structural basis for the uncoating process and acid sensitivity of EV-D68 and related viruses.

## Results and Discussion

### Current EV-D68 Strains are Acid Sensitive

The effect of acid treatment on virus infectivity was examined using a plaque assay. It was found that EV-A71 is acid resistant and retained infectivity at low pH (pH 4-6) (**Fig. 1**). In contrast, the tested EV-D68 strains are acid labile, including the prototype Fermon strain and two strains (US/MO/14-18947 and US/KY/14-18953) from the 2014 outbreak. Hereafter, US/MO/14-18947 and US/KY/14-18953 are referred to as MO and KY, respectively. Among these strains, strain MO is the most sensitive to acid (**Fig. 1**).

**Fig. 1.**
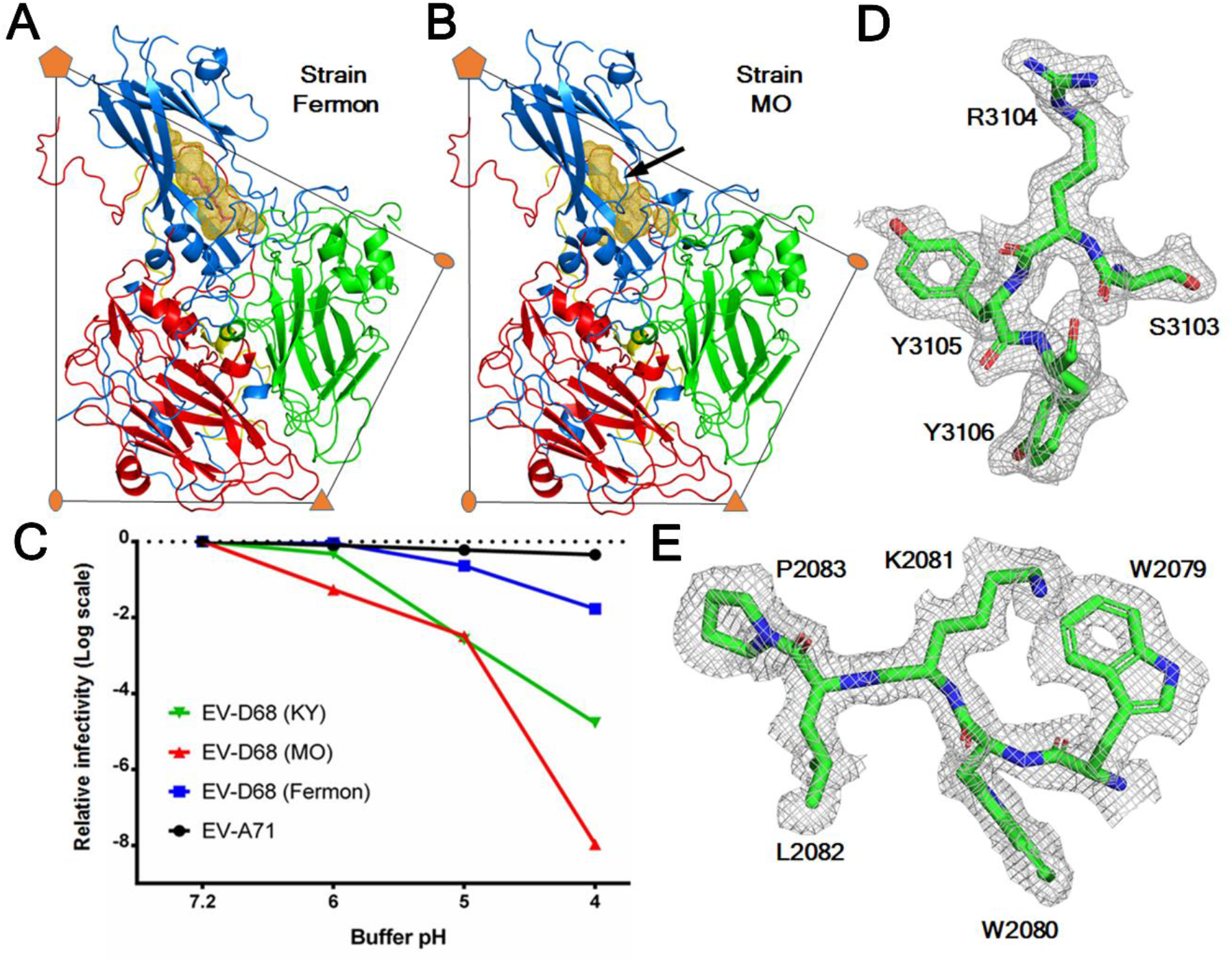
The structure of the acid sensitive EV-D68 strain MO. ***(A-B)***One protomer of the EVD68 capsid is colored blue (VP1), green (VP2), red (VP3) and yellow (VP4). The volume of the VP1 hydrophobic pocket is colored gold. A pocket factor (magenta) is present in strain Fermon***(A)*** but is absent in strain MO ***(B)***. A black arrow indicates where the collapse of the VP1 pocket of strain MO occurs. ***(C)*** Plot of changes of viral infectivity in logarithm scale as a function of buffer pH. The viruses were treated using buffer with a series of pH values and assayed to determine viral titers. ***(D-E)***. Typical map densities of strain MO at 2.2 Å resolution with the fitted atomic model.

Cryo-electron micrographs showed that there were more than 95% of full particles in a purified sample of strain MO (Prep A), which was prepared employing two rounds of density gradient centrifugation (Materials and Methods). The virus structure (dataset A_Native) was determined at 2.2 Å resolution using 16021 particles (**Fig. 1, Fig. S1, and Table S1**). In many EV structures, a hydrophobic pocket within the VP1 jelly roll accommodates a fatty acid like molecule (the “pocket factor”) that regulates viral stability (37-39). In strain MO, the pocket factor was found to be absent from the VP1 hydrophobic pocket, which is partially collapsed (**Fig. 1**). Compared with strain Fermon, the structure of strain MO shows that Ile1217 (numbering is based on the amino acid sequence of strain Fermon throughout this paper) in the VP1 GH loop moves into the pocket with the Cα atom shifted by 2.0 Å. Such a shift would cause clashes with a pocket factor were this factor bound in the pocket (**Fig. S1**).

### Acid Induces EV-D68 Uncoating

Consistent with the low stability of strain MO, a virus sample (Prep B), which was not as intensively purified as Prep A, yielded a mixture of full and empty particles within one fraction of about 0.5 ml after density gradient centrifugation (**Fig. S2*A***). To study acid triggered structural changes of the virus, Prep B was treated with either a pH 5.5 buffer (dataset B_RT_Acid) or a neutral pH buffer (dataset B_RT_Neu) at room temperature for 20 min. Two-dimensional (2D) classification of particle images (dataset B_RT_Neu) showed the presence of full and empty particles with a ratio of about 2.0:1 (full:empty) (**Fig. S2*B***). In contrast, 2D class averages of particle images in dataset B_RT_Acid showed the presence of empty particles and a particle form that contains the genome but exhibits a thinner capsid shell than native full virions (**Fig. S2*C* and *D***). The ratio between the new form of particles and empty particles was about 1.7:1, suggesting that the acid treatment had mostly induced the conversion of native full virions to the new particle form. Icosahedral reconstructions of the new form of particles (3,708 particles) and empty particles (2,150 particles) in dataset B_RT_Acid were determined to 3.3 Å and 3.8 Å resolution, respectively (**Table S1 and Fig S3**). These two forms of particles are both expanded by about 11 Å in diameter relative to native full virions. They also show significant different capsid structures than native full virions, with root-mean-square deviation (rmsd) of 4.6 Å (empty particles) and 5.7 Å (new form of particles) when aligning icosahedral symmetry axes (**Table S2**). The rmsd between any two structures was calculated based on aligning equivalent Cα atoms unless otherwise specified. Moreover, the VP1 N-terminal residues 1001-1041 and a majority of VP4 residues, which are well ordered in the map of native full virions, become disordered (or missing) in the map of the new particle form. Thus, the new particle form represents the A-particle, a proposed uncoating intermediate known to exist in other EVs (25, 26, 34). The capsid structure of empty particles resembles A-particles with an rmsd of 0.8 Å when aligning icosahedral symmetry axes (**Table S2**), as has been previously reported (26). Thus, these empty particles represent emptied particles that are formed after native full virions have released the viral genome. The emptied particles are distinct from VP0 containing native empty particles that have nearly the same capsid structure as native full virions (40, 41).

In order to mimic the environment for virus uncoating in host cells, Prep B was treated with a pH 5.5 (late endosomal pH) buffer at physiological temperature (33°C) for 20 min (dataset B_33_Acid). Similar to the aforementioned observation in the case of room temperature incubation, A-particles and emptied particles were present in dataset B_33_Acid. The cryo-EM structures of A-particles (23,082 particles) and emptied particles (19,325 particles) were determined to 2.7 Å and 2.9 Å resolution, respectively (**Table S1 and Fig. S4**). When full native virions were converted into A-particles, a VP2 helix (residues 2091-2098) and its counterpart in an icosahedral two-fold related VP2 molecule were shifted away from this two-fold axis, opening up roughly rectangularly shaped pores around the two-fold axes on A-particles (with a pore size of about 9Å × 18Å) (**Fig. 2**). Similar pores were observed on emptied particles (with a size of about 8Å × 29Å). These pores might function as sites where the genomic RNA exits, partially because a single-stranded RNA, assuming no secondary structures, has a size of slightly less than 8Å × 10Å when looking in the direction normal to the planar aromatic bases of the RNA. More importantly, the VP1 N-terminal residues 1042-1051 reside in the capsid interior of native full virions. In contrast, these residues are displaced by an rmsd of 23.6 Å in A-particles such that residues 1044-1051 traverse the capsid shell and that residue 1042 lies on the particle outer surface (**Fig. 2**). The VP3 GH loop, residues 3170 – 3188 excluding the disordered residues 3178-3183, at the particle exterior are rearranged (rmsd = 9.6 Å) to adopt an extended conformation and interact with the VP1 N-terminal residues (**Fig. 2**). These changes result in the externalization of the VP1 amphipathic helix (disordered in the A-particle structure) through a pore around the quasi-three-fold axis at the base of the “canyon” (18). The VP1 amphipathic helix (about 25 amino acids at the N-terminus) was previously shown to insert into host cell membranes (21, 42), as do also VP4 molecules (29). Therefore, these acid-induced structural changes facilitate EV-D68 uncoating.

**Fig. 2.**
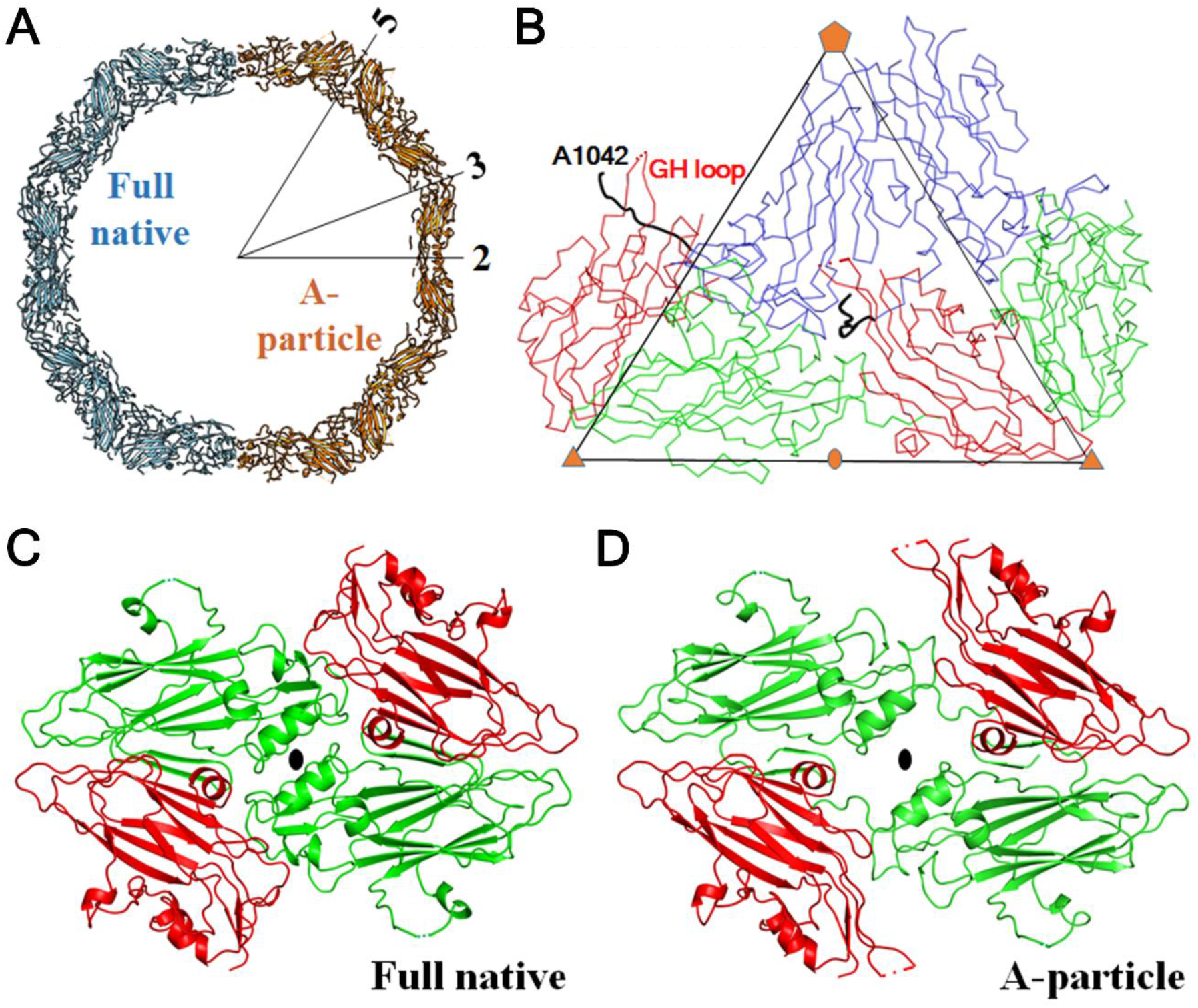
Acid-induced structural rearrangements of EV-D68 facilitate virus uncoating. ***(A)*** Structural comparison of the full native virion (pH 7.2) and A-particle (formed at pH 5.5). For each of these two structural states, a half capsid is represented as a slab of about 20 Å thick. ***(B)*** The Cα backbone representation of two neighboring protomers of the A-particle structure is colored blue (VP1), green (VP2), and red (VP3). The VP1 N-terminal residues 1042-1052, which are externalized through a quasi-three fold axis, are highlighted in black. Pores are formed around the icosahedral two-fold axes in full native virions ***(C)*** when compared with A-particles***(D)***.

### Identification of Multiple Structural Intermediates

At neutral pH, Prep B contains a heterogeneous particle population as mentioned above. To analyze the sample heterogeneity at neutral pH in detail, 2D classification and subsequently three-dimensional (3D) classification of particle images (dataset B_4_Neu) resulted in six different structural states (**Fig. S5**). These structures were determined at 3.2-3.3 Å resolution (**Table S3 and Fig. S6**). They differ mostly in particle size and in internal regions, including the VP1 N-terminal residues 1001-1053 as well as VP4. The two predominant states are full native virions (52% of all particles), and emptied particles (20% of all particles) (**Table S3**). This observation suggests that a portion of native full virions might have uncoated to produce emptied particles during virus preparation. Consistent with this prediction, two uncoating intermediates have also been identified from the whole particle population. One intermediate was found to be A-particles (about 9%), whereas the other represents a previously undescribed structural state (about 5%), named here “expanded 1 (E1) particle”. Unlike A-particles, the VP1 N-terminal residues 1001-1053 (excluding the disordered residues 1016-1019) and VP4 residues 4030-4057 are ordered in E1 particles, as is the case for full native virions (**Fig. 3**). Nevertheless, E1 particles are expanded by about 8 Å in diameter with respect to native full virions. The rmsd between these two structures when aligning icosahedral symmetry axes is 4.4 Å, whereas the rmsd between the structures of A-particles and E1 particles is 3.3 Å (**Table S4**). Thus, three distinct structural intermediates (E1, A-, and emptied particles) are involved in EV-D68 uncoating (Fig 3).

**Fig. 3.**
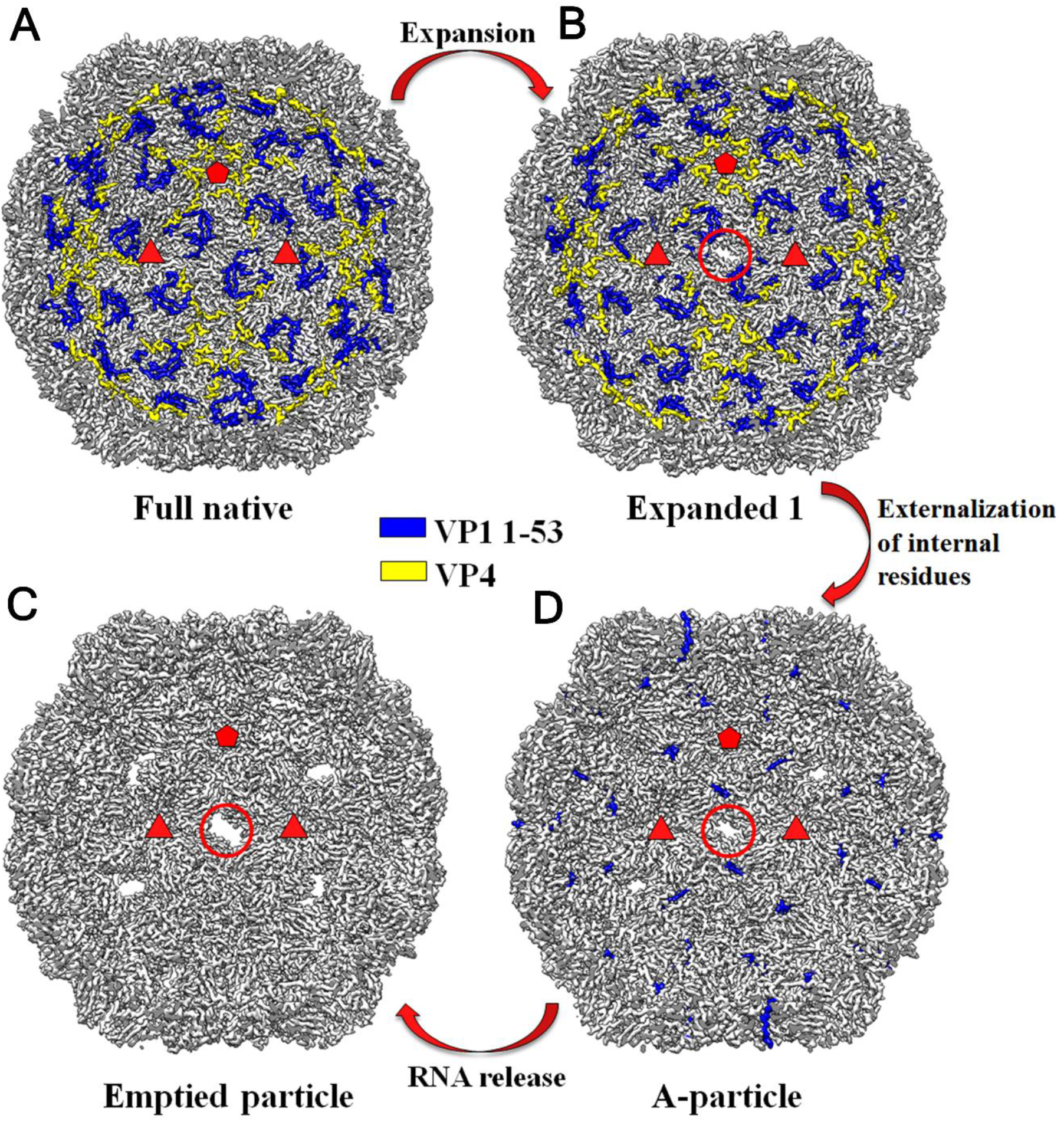
EV-D68 uncoating proceeds via multiple distinct structural intermediates. A cut-way view of each of four structural states of the EV-D68 capsid during virus uncoating. Shown are the full native virion ***(A)***, expanded 1 particle ***(B)***, A-particle ***(C)***, and emptied particle ***(D)*** when looking into the particle along an icosahedral two-fold axis. The ordered regions of VP1 N-terminal residues 1001-1053 are colored blue. The ordered regions of VP4 are colored yellow. Two red triangles and one red pentangle outline the limit of an icosahedral asymmetric unit. A red circle indicates the presence of a pore around the icosahedral two-fold axis.

In addition, two other structural states, each of which accounted for about 7% of all particles, were identified (**Fig. S7*A***). The rmsd between the full native virion and each of these two states was found to be 1.0 - 1.1 Å (**Table S4**). These two states either lacked inner densities or had rod-like densities in the particle interior (**Fig. S7*B***). These states might be abortive products during virus assembly.

### Conformational Changes of EV-D68 During Uncoating

The identification of multiple structural states of the capsid suggests that EV-D68 uncoating proceeds via a series of conformational changes. It is reasonable to assume that virus uncoating is initiated by particle expansion of full native virions to form E1 particles, producing pores on the capsid. Subsequent loss of VP4 through these pores and externalization of the VP1 N-termini result in A-particles. Ultimately the genomic RNA is released to yield emptied particles (**Fig. 3**).

Particle expansion of full native virions to E1 particles is achieved primarily through rigid body movements of capsid proteins. The centers of mass of VP1, VP2, VP3, and VP4 are translated away from the particle center by 4.5 Å, 3.4 Å, 3.9 Å, and 4.7 Å, respectively, while these proteins are rotated by 2.8°, 3.3°, 2.8°, and 3.3°, respectively. This yields a structure which has an rmsd of 1.1 Å (VP1), 1.3 Å (VP2), 1.4 Å (VP3), and 0.3 Å (VP4) when compared to the E1 particle (**Table S5**). During particle expansion the buried surface areas between VPs within a protomer (defined as VP1, VP2, VP3, and VP4) stay roughly unchanged (**Table S6**). This observation provides the basis for superimposing one protomer of the full native virion structure with a protomer in the E1 particle structure. VP2, VP3, VP4, and the VP1 regions distant from the five-fold axis are well superimposable (rmsd = 1.3 Å), whereas the VP1 β-barrel and loops near the five-fold axis are rearranged in a hinge-like manner with a shift of 0.9 Å and a rotation of 3.0°.

Pores with a size of about 6Å × 15Å are created around the two-fold axes in E1 particles, as a result of the displacement of the VP2 residues 2091-2098 near the two-fold axis with respect to full native virions (**Fig. 4**). These changes impair the interactions between neighboring pentamers. More importantly, two β-strands at the VP2 N-terminus (residues 2013-2026) in one pentamer, together with a β-strand at the VP1 N-terminus (residues 1017-1020) and the VP3 β-strands C, H, E, F in another pentamer, form a seven-stranded interpentamer β-sheet that spans from the capsid outer surface to the inner surface in full native virions (**Fig. S8*A***). This sheet and its symmetry-related equivalents help hold adjacent pentamers together and provide structural stability to the VP1 N-termini in the capsid interior (38, 43). Particle expansion disrupts this interpentamer β-sheet and weakens the interpentamer contact in E1 particles (**Figs. S8*B* and S9**). These effects are due primarily to rearrangements of VP2 residues 2012-2017 (rmsd = 4.0 Å), VP2 residues 2026-2030 (rmsd = 4.3 Å), and VP1 residues 1020-1026 (rmsd = 8.8 Å) with respect to full native virions (**Fig. S10**). Furthermore, the VP3 GH loop in E1 particles becomes partially disordered and moves outwards (away from the virus center) with an rmsd of 5.6 Å (residues 3180 and 3186-3188) (**Figs. S10 and S11**). These changes might precede pore opening at the base of the canyon, similar to what was previously proposed (44). Thus, structural alterations from full native virions to E1 particles not only lead to impaired pentamer-pentamer interactions (**Table S7**), but also prime the exiting of VP4 and the externalization of the VP1 N-termini.

**Fig. 4.**
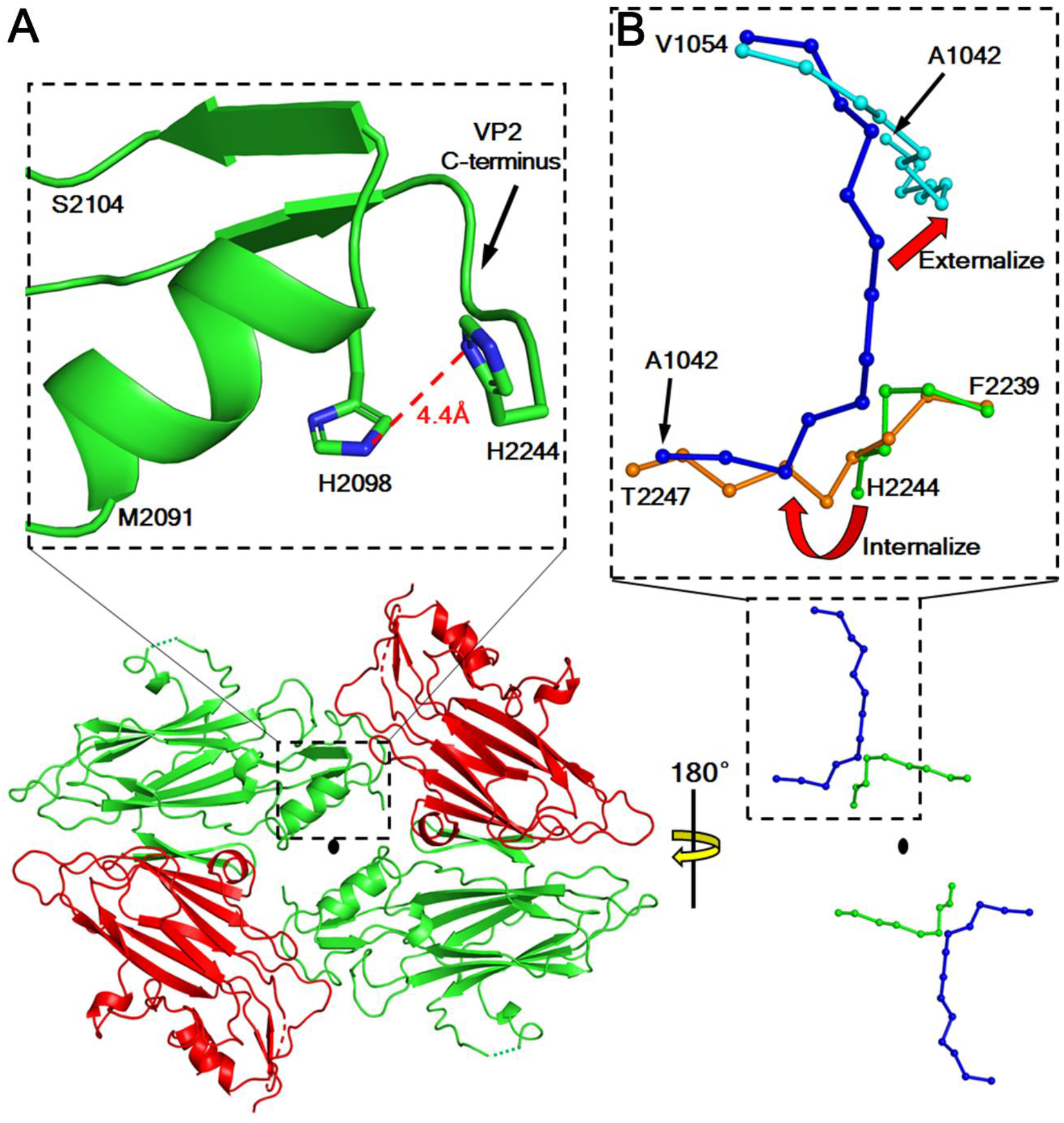
A histidine-histidine pair near the two-fold axes facilitates the formation of A-particles under acidic conditions. **(*A*)** In the lower part of the panel, pores are open around the two-fold axes in E1 particles. VP2 and VP3 are colored green and red. The enlarged portion in the upper part of the panel shows the locations of the histidine pair, His2098 and His2244. The smallest distance between the imidazole moieties of these two histidines is 3.8 Å. ***(B)*** Superposition of equivalent protomers of expanded E1 particles (VP1: blue, VP2: green) and A-particles (VP1:cyan; VP2: orange). When E1 particles are converted into A-particles, internalization of the VP2 C-terminal tails promotes the externalization of the VP1 N-termini.

Superposition of equivalent protomers in E1 particles and A-particles shows that VP2, VP3, and the five-fold distant regions of VP1 are well aligned with an rmsd of 1.1 Å. However, the VP1 β-barrel and five-fold proximal loops undergo hinge-like motions with a rotation of 1.9°. The VP2 C-terminal tail (residue 2242 to the carboxyl end) at the capsid exterior in E1 particles becomes internalized in A-particles. This generates enlarged pores around the two-fold axes, facilitating the exiting of VP4 molecules (45). Moreover, the internalized VP2 C-terminal tail would clash with the N-terminal residues 1042-1044 of VP1 in a neighboring, five-fold related protomer if the VP1 N-terminus were to stay stationary (**Fig. 4**). Structural reorganization of the VP3 GH loop and VP1 GH loop in A-particles with rmsd values of 11.5 Å (residues 3170-3178) and 2.7 Å (residues 1202-1207) with respect to E1 particles creates room near the quasi-three-fold axes (**Figs.S11 and S12**). Collectively, these conformational changes from E1 particles to A-particles promote the externalization of VP1 N-termini through holes at the base of the canyon (**Fig. 3 and Fig. S11**).

The final stage of uncoating involves genome release from a specific pore around a twofold axis in A-particles, generating emptied particles (46, 47). Given the high structural similarity between these two states as mentioned above, structural analyses make it difficult to identify the trigger that initiates RNA release. Nevertheless, this process has been reported to depend on interactions of A-particles with host cell membranes (30, 42, 48), disruption of the secondary structures of the viral RNA (49), and electrostatic repulsion between the negatively-charged RNA and negatively-charged residues lining the inner capsid surface (26).

### Molecular Basis for Acid Lability

Unlike members of the EV-A, EV-B and EV-C species that have a well-formed pocket factor with a long aliphatic chain, the VP1 hydrophobic pockets of RVs and EV-D68 either contain a pocket factor with a short aliphatic chain (RV-A2, RV-A16, EV-D68 strain Fermon) or cannot accommodate a pocket factor at all due to the pocket being collapsed as in RV-B3, RV-B14, and EV-D68 strain MO (36). Furthermore, RV-A2, RV-A16, and RV-C15 have particularly small interaction areas at the pentamer-pentamer binding interface (**Table S7**). In comparison with other EVs, the above mentioned structural features of EV-D68 and RVs would lead to enhanced conformational fluctuations of the capsid (50) and promote irreversible conformational changes from the full native virion to an E1 particle. As mentioned above, the E1 particle state primes the externalization of the VP1 N-termini and loss of VP4, which are major features of the A-particle state. Low pH conditions probably facilitate the conversion into A-particles in three ways:

i. Particle expansion from full native virions to E1 particles could lead to influx of protons through transient openings around the five-fold axes, causing conformational changes of fivefold proximal regions at the capsid interior, including VP4. Similar changes that precede the release of VP4 have been reported for RV-B14 under acidic conditions (51).
ii. Protons could enter into the capsid through pores around the two-fold axes and disrupt polar interactions that stabilize the VP1 N-termini, which are already destabilized in E1 particles relative to full native virions. Acid is also known to modulate the conformational states of the GH loops in VP1 and VP3 around the quasi-three-fold axes (51, 52). These acid-induced changes would probably drive the externalization of the VP1 N-termini.
iii. The E1 particle state shows partially disrupted seven-stranded interpentamer β-sheets because of rearranged VP1 and VP2 N-termini. Structural rearrangements of these regions have been proposed to regulate the accessibility of interpentameric histidine residues to the acidic environment (53). This may further impair the interactions at the pentamer-pentamer interface, as is observed in A-particles with respect to E1 particles (**Table S7**).

Differences of amino acid sequences in VP1, VP2, VP3 and VP4 might explain why the two recent EV-D68 strains, MO and KY, are more sensitive to acid than the prototype Fermon strain. The major difference between strain Fermon and strains MO/KY involves substitutions of 15 ionizable amino acid residues on the virus outer surface (**Table S8**). Among these residues, strains MO and KY have a His at position 2098, whereas strain Fermon has a Tyr. The E1 particle structure of strain MO shows a pair of histidine residues (His2098 and His2244 from the same VP2 molecule) that line the pores around two-fold axes. The imidazole moieties of these residues are accessible to solvent and have a distance of about 4 Å from each other. The side chain of histidine has a pKa of about 6. Thus, it is probable that protonation of these histidines under acidic conditions (e.g., pH 5.5) results in electrostatic repulsion that drives the internalization of the VP2 C-terminal tails and promotes the externalization of the VP1 N-termini (**Fig. 4**). Thus low pH can facilitate the formation of the A-particle state. The poor infectivity of A-particles (54) might account for why acid treatments lead to loss of EV-D68 infectivity (**Fig. 1**).

Sequence alignments show that His2244 is completely conserved among 469 EV-D68 strains, whereas at position 2098 the histidine is replaced by tyrosine in 31 strains. These 31 strains were isolated between 1962 and 2013 (**Table S9**). Thus strains that have a His at position 2098 include those from the 2014 outbreak in the United States (55) as well as those isolated from patients who developed acute flaccid myelitis (56). These current strains might have low stability under acidic conditions, a property that probably allows efficient virus uncoating within host cells.

### Implications for Cell Entry of EV-D68

Previous reports have identified sialic acid (a carbohydrate moiety) and intercellular adhesion molecule 5 (ICAM-5), a sialic acid containing glycoprotein, as cellular receptors for EV-D68 (11, 14-16). The present work indicates that endosomal acidification might serve as a trigger for EV-D68 uncoating in host cells. Sialic acid receptor binding to the Fermon strain has previously been shown to cause ejection of the pocket factor that destabilizes the virus (14), resulting in a virus that is much like strain MO because both structures show the absence of a pocket factor. The infectivity of the Fermon strain is slighted impaired at pH 5-6 (**Fig. 1**), suggesting that endosomal acidification alone is probably insufficient for uncoating of this strain. This observation raises the possibility that sialic acid receptor binding increases the pH threshold for inducing EV-D68 uncoating. Likewise, strain KY uncoats after incubation with soluble ICAM-5 at pH 6.0, whereas the uncoating process is less efficient when the virus was incubated without a receptor molecules at pH 6.0 (16). Hence, cellular receptors (e.g, sialic acid or ICAM-5) and endosomal acidification probably assist EV-D68 entry into host cells in a synergistic manner, as has previously been proposed for other EVs (57, 58). In this way, receptor binding can prime EV-D68 uncoating, which then occurs in intracellular compartments that have a suitable environment (e.g., acidic pH) for genome release. Nevertheless, the synergistic action of cellular receptors and low pH might depend on virus strains. For instance, the infectivity of strain MO is reduced by about 100-fold at pH 6.0 (**Fig. 1**). The A-particle state of this strain is formed at pH 5.5. Thus, endosomal acidification alone is probably sufficient for triggering uncoating of strain MO. In this sense, receptor molecules might only lead to internalization of the virus via endocytosis, and also act as anchors to place the virus in close proximity to the membranes of appropriate intracellular compartments where virus uncoating can take place.

In summary, cryo-EM analyses of the acid sensitive EV-D68 have shown the involvement of multiple structural intermediates in the viral uncoating pathway. A set of high resolution structures presented here provide the basis for developing antiviral therapeutics that would interfere with structural rearrangements of EV-D68 during cell entry. Moreover, the identification of expanded E1 particles, a previously unknown intermediate, suggests that the conformational fluctuations of enterovirus capsids may account for the differences in acid lability among enteroviruses.

### Data Deposition

The atomic coordinates of A_Native-full, B_RT_Acid-Aparticle, B_RT_Acid-Emptied, B_33_Acid-Aparticle, B_33_Acid-Emptied, B_4_Neu-Full-Native, B_4_Neu-E1, B_4_Neu-Aparticle, B_4_Neu-Emptied, B_4_Neu-Abortive1, and B_4_Neu-Abortive2 have been deposited with the Protein Data Bank (accession numbers 6CSG, 6CS6, 6CSA, 6CS4, 6CSH, 6CRR, 6CS3, 6CRS, 6CRU, 6CRP, 6CS5). The cryo-EM maps of A_Native-full, B_RT_Acid-Aparticle, B_RT_Acid-Emptied, B_33_Acid-Aparticle, B_33_Acid-Emptied, B_4_Neu-Full-Native, B_4_Neu-E1, B_4_Neu-Aparticle, B_4_Neu-Emptied, B_4_Neu-Abortive1, and B_4_Neu-Abortive2 have been deposited with the Electron Microscopy Data Bank (accession numbers EMD-7599, EMD-7593, EMD-7598, EMD-7589, EMD-7600, EMD-7569, EMD-7583, EMD-7571, EMD-7572, EMD-7567, EMD-7592).

## Materials and Methods

### Viruses

EV-A71 (strain MY104-9-SAR-97, GenBank ABC69262.1) was provided by Jane Cardosa (Universiti Malaysia Sarawak, Malaysia). EV-D68 strain Fermon CA62-1 (strain Fermon, GenBank: AY426531.1) was provided by M. Steven Obserste (Centers for Disease Control and Prevention of the United States). Two EV-D68 strains, US/MO/14-18947 (strain MO, GenBank: AIS73051.1) and US/KY/14-18953 (strain KY, GenBank: AIS73057.1), from the 2014 outbreak in the United states were obtained from BEI Resources, National Institute of Allergy and Infectious Diseases, National Institute of Health. All viruses were propagated in human rhabdomyosarcoma cells (RD, ATCC CCL-136) and stored at −80 °C.

### Virus Growth and Purification

A sample (Prep A) for structure determination of the full EV-D68 virion (strain MO) was prepared in the following way (36). Briefly, RD cells were infected with EV-D68 at a multiplicity of infection of about 0.01. Cells and supernatant were harvested at three days post infection and were then separated by centrifugation. Cell pellets were subjected to multiple cycles of freezing and thawing followed by centrifugation to remove cell debris. The resultant supernatant was combined with the original supernatant and used for ultracentrifugation at 278,000 × g (Ti 50.2 rotor) for 2h at 4°C. The resultant pellets were resuspended in buffer A (250 mM HEPES, 250 mM NaCl, pH 7.5) and treated sequentially with 5 mM (final concentration throughout the treatments) MgCl2, 10 µg/ml DNase, 7.5 mg/ml RNase, 0.8 mg/ml trypsin, 15mM EDTA, and 1% (w/v) sodium n-lauryl-sarcosinate. The crude virus sample was sedimented through a potassium tartrate density gradient (10% - 40% (w/v)). A band in the middle of the tube was extracted and subjected to buffer exchange. The resultant sample was further purified using an iodixanol (OptiPrep, Sigma) density gradient (10% - 51% (v/v)) at 175,000 × g (SW 41 Ti rotor) for 2h at 4°C. Electron micrographs of the final sample verified the presence of more than 95% of full particles (**Fig. S1A)**.

A virus preparation (Prep B) that contained a heterogeneous population of EV-D68 particles was used for structural studies at both neutral pH and acidic pH. Procedures for virus infection, sample collection, initial centrifugation, and treatment of cell pellets followed the same procedure as described above. Subsequently, polyethylene glycol 8000 (PEG8000) (40% w/v stock solution) and NaCl (powder) were added into the original supernatants (after infection) to reach a final concentration of 8% PEG8000 and about 500 mM of NaCl. After low speed agitation at 4°C for about 6h, the mixture was spun down. The resultant pellets were resuspended in buffer A, which was combined with the previous supernatant from the step that dealt with the cell pellets. The remaining steps were the same as mentioned above, except that the crude virus sample was purified through only one round of density gradient centrifugation using the iodixanol gradient.

### Acid Sensitivity Assay

Four different viruses were used, including the acid resistant EV-A71, the acid labile EV-D68 strain Fermon, and two EV-D68 isolates (strains MO and KY) from the 2014 outbreak in the United States. Purified viruses were treated with phosphate-citrate buffer (100 mM Na_2_HPO_4_ and 50 mM citric acid) at pH 4.0, 5.0, 6.0, or 7.2 at 33°C for about 40 min. The resultant samples were neutralized back to pH 7.1-7.2 using 400 mM Na_2_HPO_4_ and 200 mM citric acid (pH 7.3) before being assayed for determination of viral titers.

### Cryo Electron Microscopy

About 2.8 µl of sample was applied onto a 400 mesh continuous carbon grid (Ted Pella Inc.). Immediately after blotting for about 8s, the grid was vitrified in liquid ethane that was pre-cooled by liquid nitrogen. Frozen, hydrated particles that were embedded in a thin layer of vitreous ice were imaged with a K2 Summit direct electron detector (Gatan) using a Titan Krios transmission electron microscope (FEI) operating at 300 kV. Cryo-EM data on strain MO were automatically collected using the program Leginon (59). The dose rate was kept at approximately 8e^-^/pixel/s for data collection. For structure determination of native full virions using Prep A, movies of frozen, hydrated virus particles (dataset A_Native) were collected at a nominal magnification of 22,500× in super resolution mode with defocus values ranging from 0.3 to 3.0 µm. A total electron dose of about 36 e^-^/Å^2^ was fractionated into 38 frames in every movie with a frame rate of 200ms/frame. For initial low pH studies using Prep B, viruses were treated with phosphate-citrate buffer (100 mM Na_2_HPO_4_ and 50 mM citric acid) to reach a final pH of 5.5 (dataset B_RT_Acid) or a pH of 7.2 (dataset B_RT_Neu), followed by incubation at room temperature for 20 min and, subsequently, neutralization with 400 mM Na_2_HPO_4_ and 200 mM citric acid (pH 7.8). Data were collected at a nominal magnification of 22,500× in super resolution mode. The defocus range for datasets B_RT_Acid (144 movies) and B_RT_Neu (87 movies) were 0.9-4.8 µm and 1.3-3.6 µm, respectively. For dataset B_RT_Acid, a total electron dose of about 28 e^-^/Å^2^ was fractionated into 30 frames (200ms/frame). For dataset B_RT_Neu, a total electron dose of about 25 e^-^/Å^2^ was fractionated into 27 frames (200ms/frame). For low pH studies using Prep B to mimic conditions during virus infection, viruses were treated at pH 5.5 similarly to the aforementioned procedure except that the incubation temperature was changed to 33°C. Data (dataset B_33_Acid) were collected at a nominal magnification of 22,500× with defocus values ranging from 0.5 to 3.5 µm. A total electron dose of about 38 e^-^/Å^2^ was fractionated into 40 frames (200ms/frame). For analyzing the heterogeneous particle population of Prep B (stored at 4°C) at neutral pH, data (dataset B_4_Neu) were collected at a nominal magnification of 18,000× in electron counting mode with defocus values ranging from 1.7 to 5.3 µm. A total electron dose of about 45 e^-^/Å^2^ was fractionated into 60 frames (250ms/frame).

### Image Processing

For all datasets, movie frames were subjected to motion correction using a modified version (Wen Jiang, Purdue University) of MotionCorr (60). The aligned frames were summed up to produce individual micrographs, which were used to estimate contrast transfer function (CTF) parameters using CTFFIND3 (61). For the datasets A_Native, B_RT_Acid, and B_RT_Neu, virus particles were picked from the micrographs manually using e2boxer.py in the EMAN2 program package (62). For all other datasets, particle selection was performed first manually using e2boxer.py and subsequently automatically using the program DogPicker (63) based on templates derived from manually selected particles. Particles were subsequently boxed and extracted from the micrographs. The process was integrated into the Appion data processing pipeline (64). The resultant particle images were subjected to two-dimensional (2D) classification using the program Relion (65), which identified and removed some low quality particles and separated images of full particles from those of empty particles.

The following reconstruction procedures were employed for all datasets except for dataset B_4_Neu using the program jspr (66). In brief, particle images (8× binned, with a pixel size of 5.20 Å/pixel) were divided into two halves. For each half, random initial orientations were assigned to individual particles, allowing for reconstruction of multiple icosahedral three-dimensional (3D) models, from which a suitable initial model was selected. The best particle orientation and center of each particle image was searched with respect to projections of the initial reference model. The reference model for the next iteration was reconstructed from particle images employing parameters for orientation and center determined in the current iteration. The refinement procedure was then extended to 4× binned and then 2× binned data. Specifically, for dataset A_Native, the procedure was extended to unbinned data. Subsequently, multiple parameters were included in the refinement process, which were parameters for particle orientation, particle center, beam tilt, defocus, scale, astigmatism, and anisotropic magnification distortion (67, 68). To achieve 3D reconstructions with the highest possible resolution, particle images (dataset B_RT_Acid only) were re-extracted from micrographs that were generated by summing aligned frames 3-16. In this way, frames that underwent large motions and that contained limited high resolution information due to radiation damage were discarded. Frames 3-16 were selected using a trial-and-error approach in which different combinations were tested, including frames 3-9, 3-16, 3-23, and 3-30. For datasets A_Native and B_33_Acid, movie frames were aligned using the program MotionCor2 (an improved version of MotionCorr) (69), in which the first frame of each movie was discarded due to large motions and high resolution information in late frames were down-weighted using a reported dose-weighting scheme (70). Micrographs were generated by summing aligned frames. Particle images were re-extracted from individual micrographs without changing the coordinates of individual particles on every micrograph and used for reconstructing the structures of full native virions, A-particles, and emptied particles at 2.17 Å, 2.73 Å, and 2.90 Å resolution, respectively. Fourier shell correlation (FSC) of two interdependently calculated half-maps (masked with a soft mask) was used to estimate the resolution of the final EM maps using an FSC cut-off of 0.143 (71, 72). The maps were sharpened by applying a negative B-factor and filtered with an FSC-curve based low pass filter (71).

The following procedures were applied to dataset B_4_Neu. After 2D classification of all particle images in the dataset, the resultant full particle images (4× binned, 6.48 Å/pixel) were used to generate initial 3D models, from which a suitable initial model was selected. The refinement process was performed using a projection matching approach as described above. After multiple iterative cycles when the process converged, the resultant 3D model was essentially an average of all possible structural states present in the collection of full particle images. The model was low pass filtered to 60 Å resolution and then utilized as a reference model for 3D classification of full particle images (4× binned) using the program Relion (65), where the number of classes was four and where icosahedral symmetry was imposed. Particle images from two of the resultant four classes yielded 3D reconstructions that were nearly identical to each other by visual inspection. Thus, particle images from the two classes were combined into one class. The same process was also applied to images of empty particles. Hence, all particle images in the dataset were classified into a total of six classes (three for full particles and three for empty particles), which represented six different structural states. Procedures for cryo-EM structure determinations were the same when using each class of particle images, as were detailed above.

### Model Building and Refinement

The same procedures were employed for all atomic structures presented in this work. The coordinates of the EV-D68 Fermon strain excluding the pocket factor and water molecules (PDB accession number 4WM8) were used as a starting atomic model. It was manually fitted into the EM map using Chimera (73). Then multiple cycles of model rebuilding in Coot (74) and real space refinement against the EM map using Phenix (75, 76) yielded an atomic model that fitted well into the map density as judged by visual inspection. A mask that contained all grid points within a radius of 5 Å around each atom of the atomic model was used to cut out a map segment from the EM map. This map segment was placed into a pseudo crystallographic unit cell with space group P1 and back-transformed into structure factors. The atomic model was subjected to refinement of atomic coordinates, B factors, and occupancy against these pseudo crystallographic structure factors in reciprocal space using REFMAC5 (77). The resultant atomic model was used for real space refinement with 60-fold non-crystallographic symmetry constraints using Phenix (75, 76). Water molecules were added in Coot (74). The final atomic models were validated according to the criteria of MolProbity (78). All figures were generated using Pymol (https://pymol.org/) or Chimera (73). Structural comparisons were done using HOMOLOGY (79). Oligomers of capsid protomers were produced using VIPERdb (80). Buried surface areas at protein-protein interacting interfaces were calculated using PISA (81).

## ACKNOWLEDGMENTS

We thank M. Steven Oberste of the Centers for Disease Control and Prevention for providing the Fermon strain of EV-D68, and Jane Cardosa of Universiti Malaysia Sarawak, Malaysia, for supplying the EV-A71 strain MY104-9-SAR-97. The following reagents were obtained through BEI Resources, NIAID, National Institutes of Health: Enterovirus D68, US/MO/14-18947 (NR-49129) and US/KY/14-18953 (NR-49132). We are grateful to Thomas Klose, Zhenguo Chen, Yingyuan Sun, Valorie Bowman, and Wen Jiang for help with cryo-EM analysis, and Sheryl Kelly for help preparing this manuscript. This work was supported by NIAID grant AI011219 to MGR.

## Author contribution

Y.L. designed the study; Y.L. and J.S. performed the experiments; Y.L. and M.G.R. analyzed data and wrote the manuscript. The authors declare no conflict of interest.

## Supplementary Information for

**Supplementary Information includes**

**Figures S1 – S12**

**Tables S1 – S9**

**Fig. S1.**
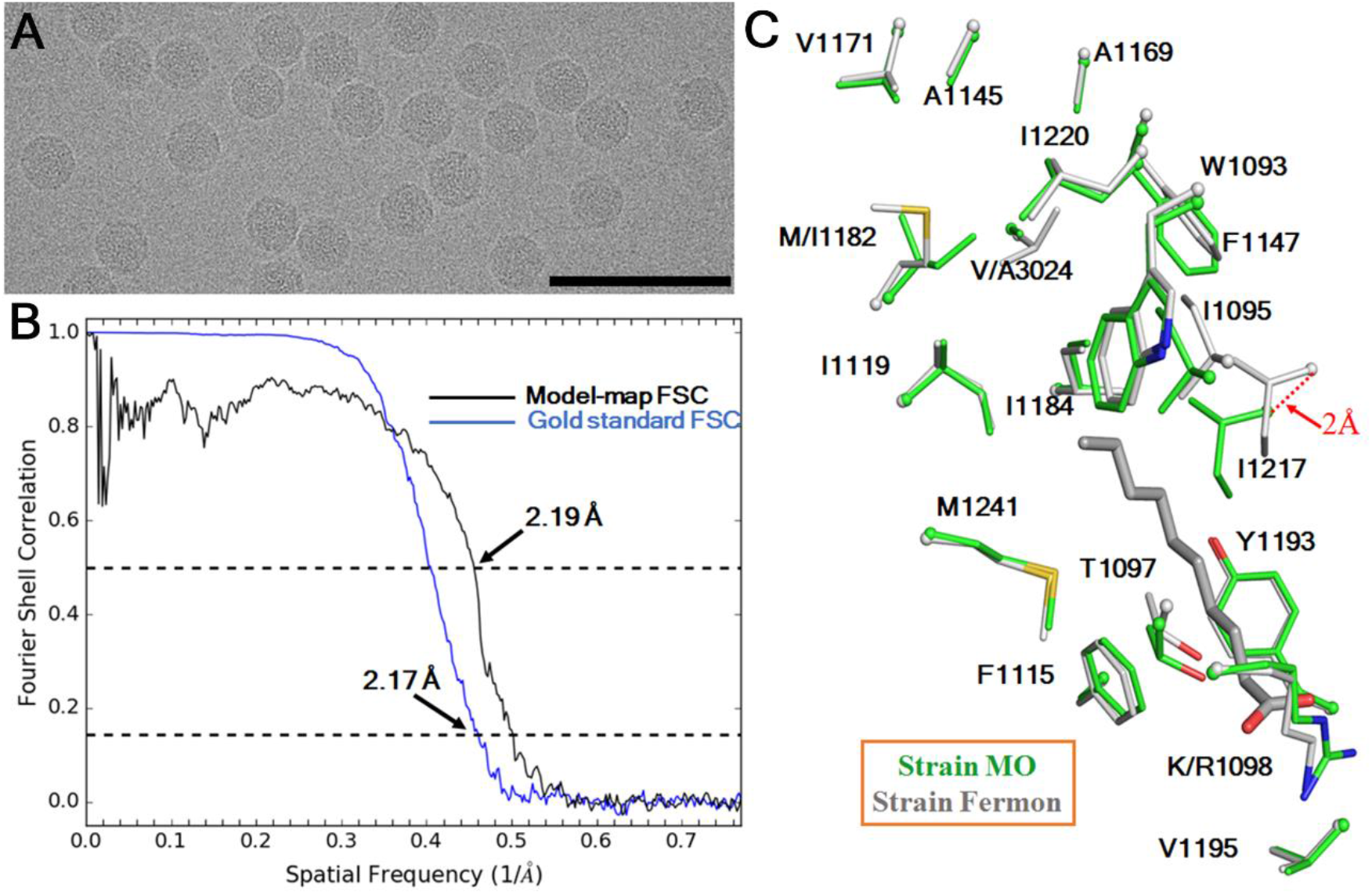
The cryo-EM structure of EV-D68 strain MO. ***(A)*** A typical portion of an electron micrograph of strain MO. This micrograph was collected at a defocus of 1.3 µm. Scale bar: 100 nm. ***(B)*** Estimation of map resolution based on Fourier shell correlation (FSC) curves. Gold standard FSC refers to the FSC between two independently reconstructed half maps using an FSC cutoff of 0.143 (71, 72). Model-map FSC refers to the FSC between the final cryo-EM map and a map calculated based on the atomic model using an FSC cutoff of 0.5 (71). ***(C)*** A close-up view of residues lining the VP1 hydrophobic pockets in strain MO (green) and in strain Fermon (grey). A pocket factor (grey) is present in strain Fermon but is absent in strain MO. In the pocket, these two strains differ mainly in two hydrophobic residues at positions 1182 and 3024. Among 469 EV-D68 strains for which the nearly complete genome sequences are available, the occurrences of Met and Ile at position 1182 are 58% and 42%, respectively. The occurrences of Ala and Val at position 3024 are 52% and 48%, respectively.

**Fig. S2.**
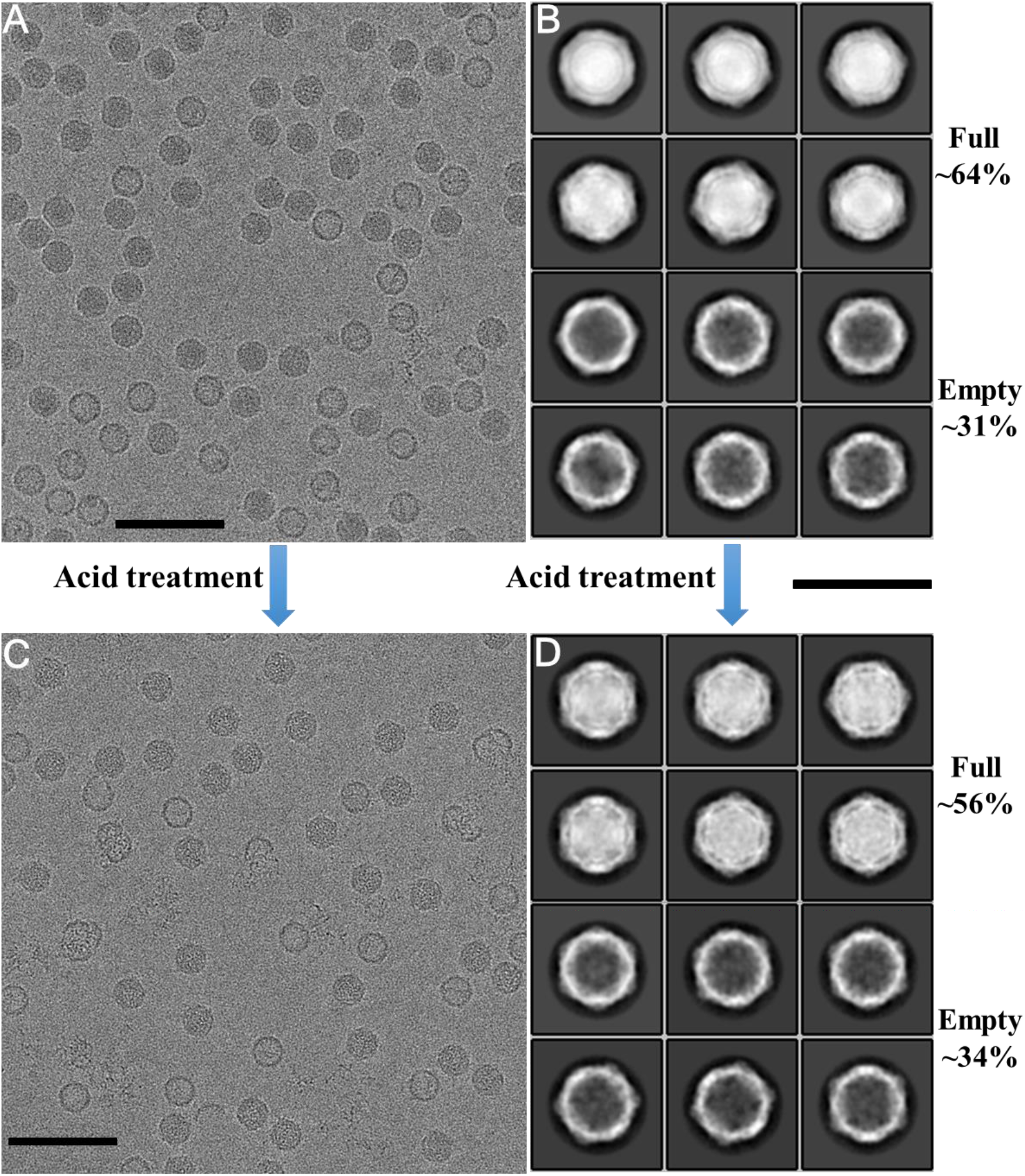
Acid induces structural alterations of EV-D68 strain MO. Typical electron micrographs show EV-D68 particles that were treated with either a pH 7.2 buffer ***(A)*** or a pH 5.5 buffer ***(C)*** at room temperature for 20 min. Scale bar: 100 nm. The corresponding 2D class averages of particle images are shown in ***(B)*** and ***(D)***, respectively. The percentage of full or empty particles among all particles is given on the right side. The scale bar for ***(B)*** and ***(D)*** represents 50 nm.

**Fig. S3.**
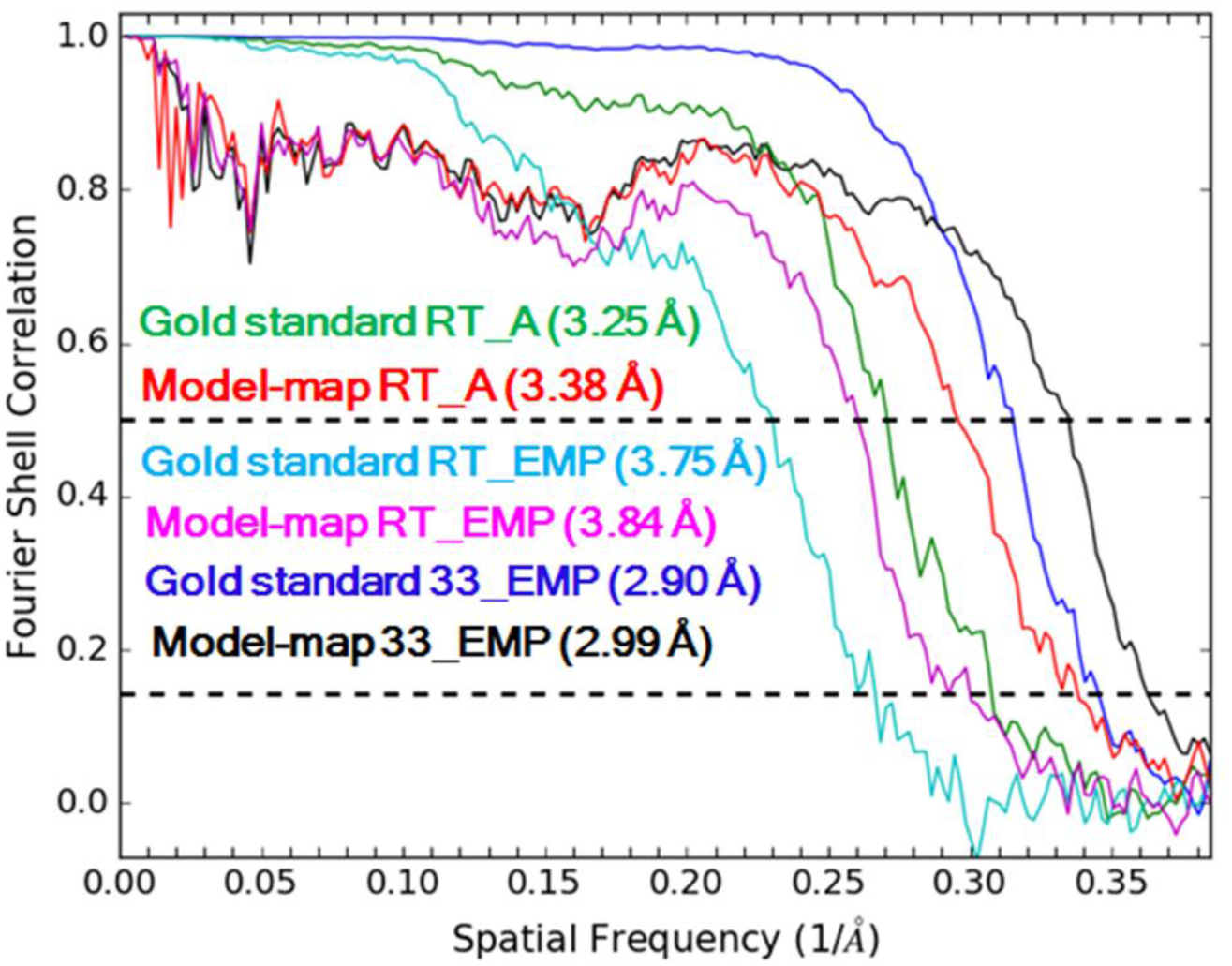
Assessment of map resolutions based on FSC curves. RT_A and RT_EMP refer to A-particles and emptied particles in dataset B_RT_Acid. 33_EMP refers to emptied particles in dataset B_33_Acid. The FSC curves are defined as in **Fig. S1**.

**Fig. S4.**
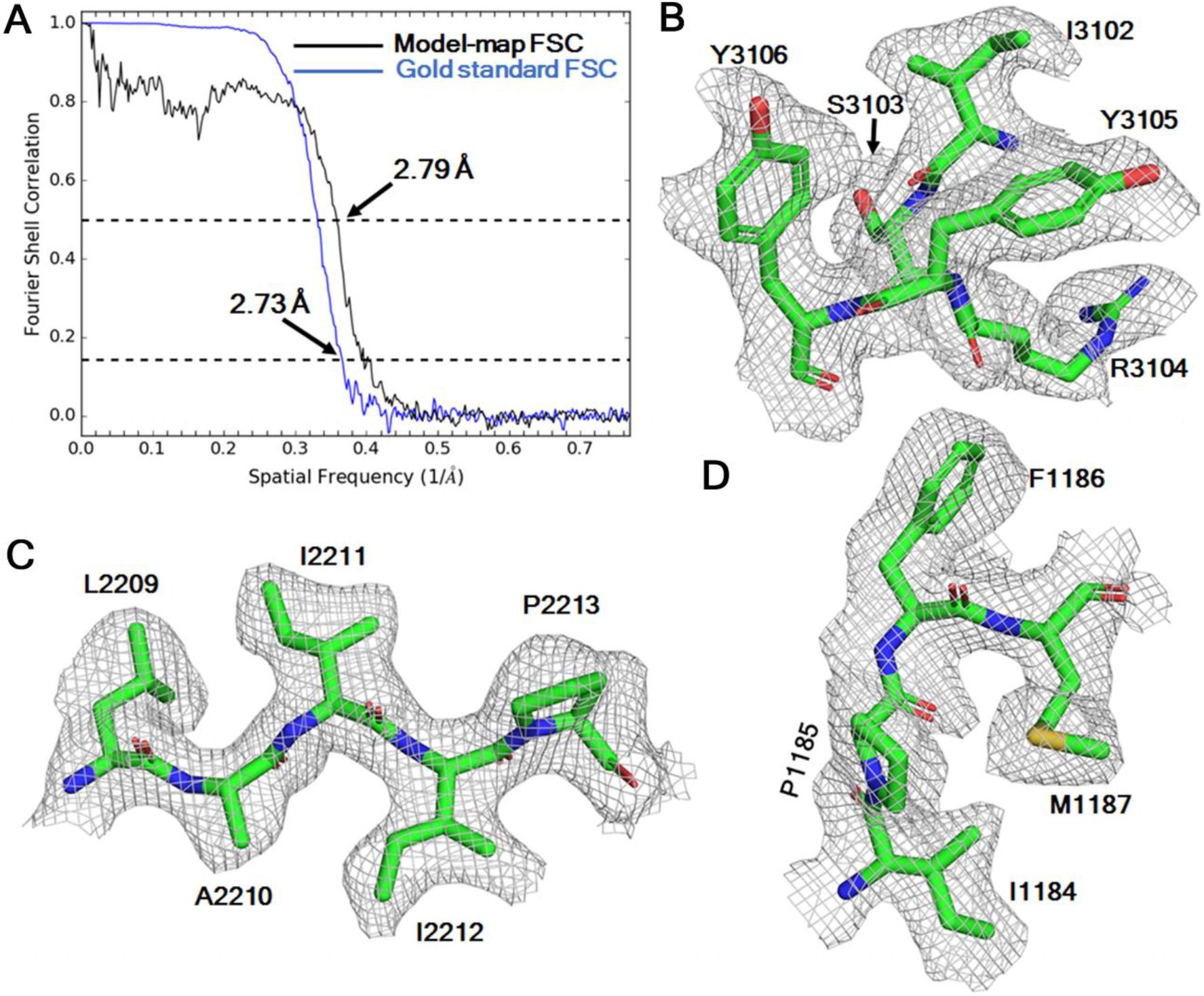
The 2.7 Å resolution structure of A-particles. ***(A)*** The resolution of the cryo-EM map was estimated based on FSC curves, defined as in **Fig. S1**. ***(B-D)*** Typical densities of the cryoEM map with the fitted atomic model.

**Fig. S5.**
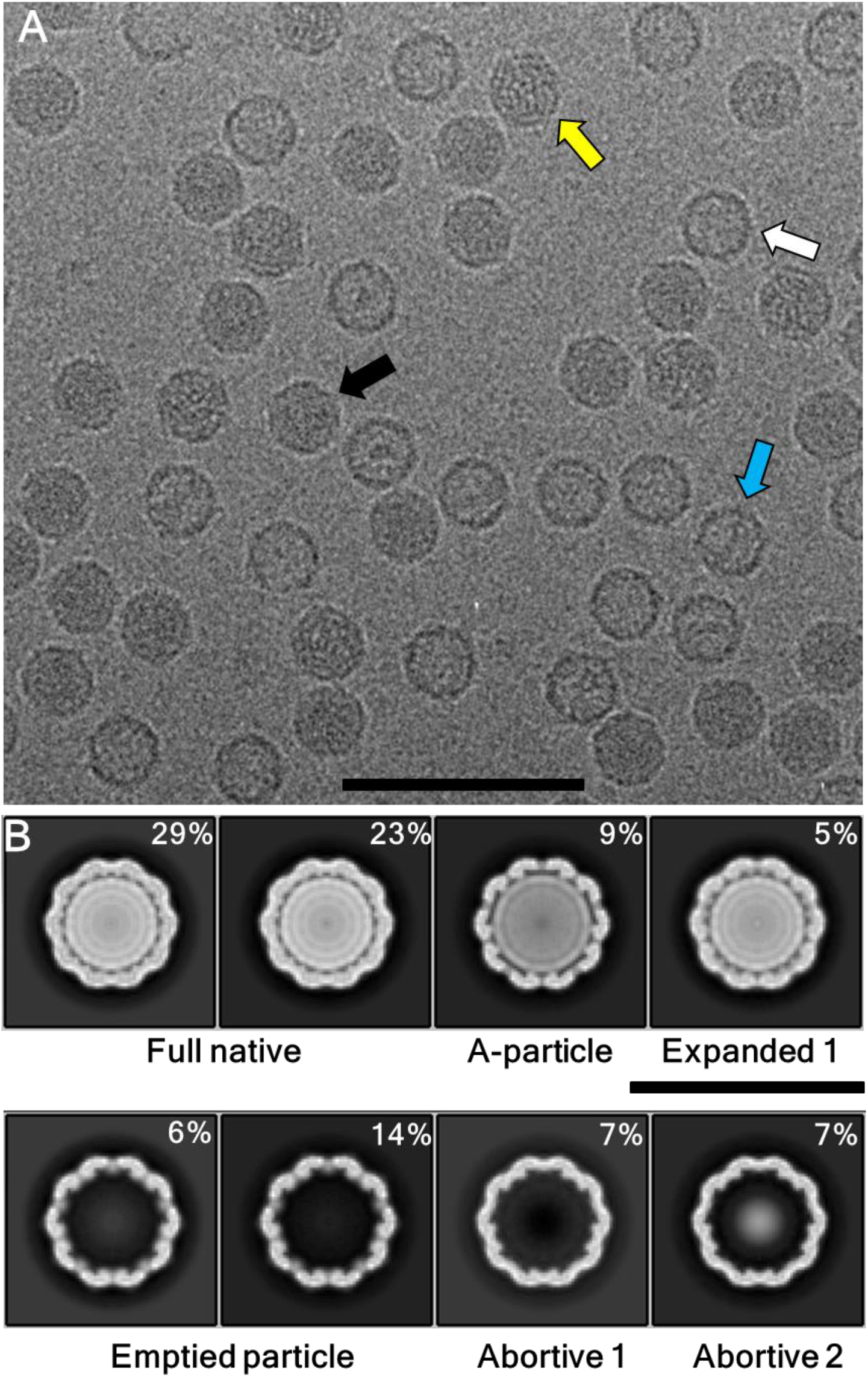
Cryo-EM analyses of a mixed population of EV-D68 particles demonstrate the presence of multiple distinct structural states. ***(A)*** A typical electron micrograph of EV-D68 particles kept at 4°C and at pH 7.2. Arrows with different colors indicate distinct particle forms judged by visual inspection. Scale bar: 100 nm. ***(B)*** Three-dimensional (3D) reconstructions obtained after 3D classification show differences between structural states. The central slice of each reconstruction is shown. The percentage of particles, which were used for each reconstruction, among all particles is given at the upper right corner. Scar bar: 50 nm.

**Fig. S6.**
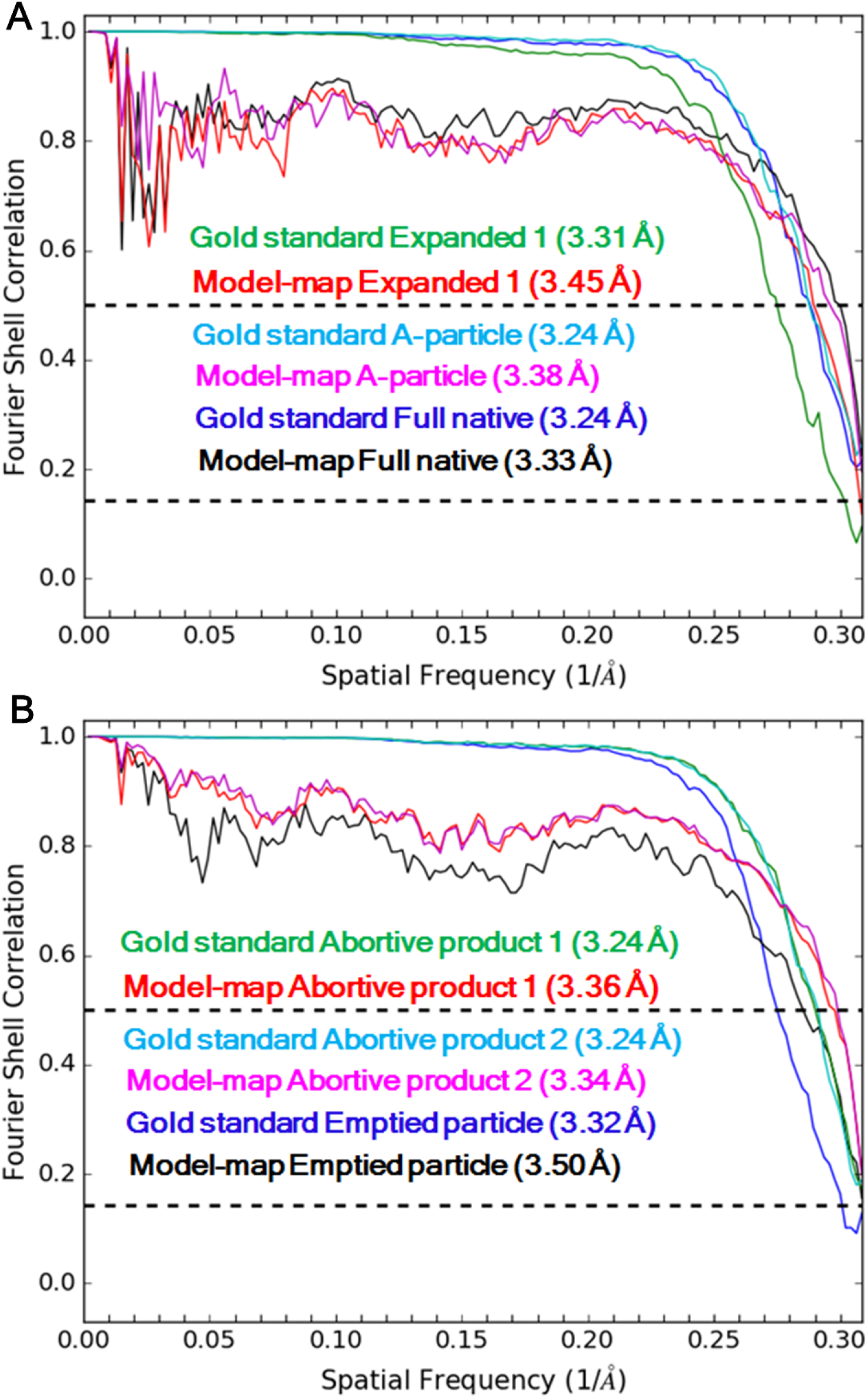
Assessment of the resolution of cryo-EM maps (dataset B_4_Neu) based on FSC curves. Shown are FSC curves for each of the three full particle states ***(A)*** and those for each of the three empty particle states **(B)**. Gold standard FSC and model-map FSC are defined as in **Fig. S1**.

**Fig. S7.**
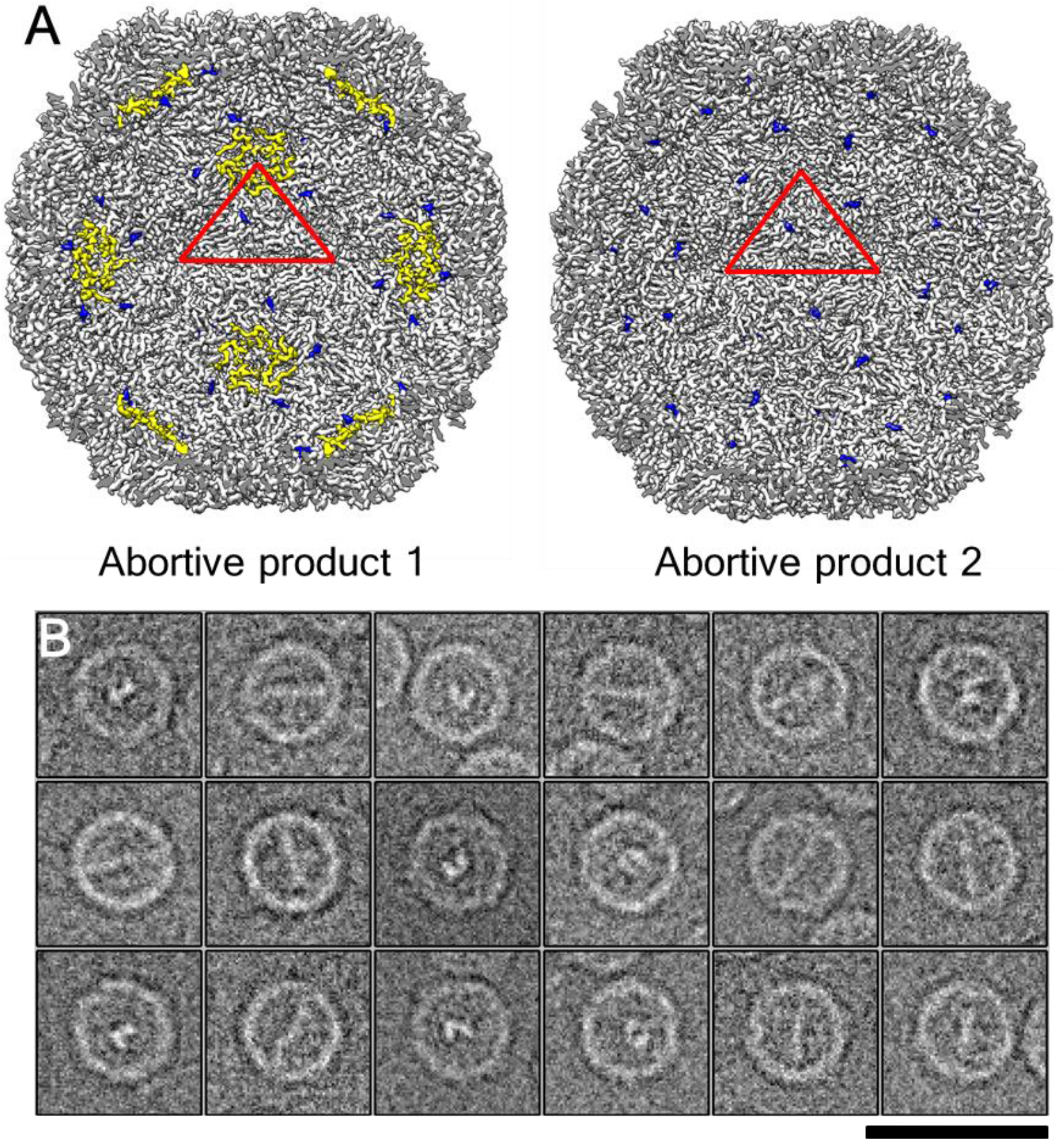
Multiple structural states of EV-D68 are present at neutral pH. ***(A)*** Cut-way views of two structural states, which probably represent abortive products during EV-D68 assembly, when looking into the particle along an icosahedral two-fold axis. A red triangle indicates an icosahedral asymmetric unit. The internal regions are colored as in **Fig. 3**. ***(B)*** Typical cryo-EM images of abortive product 2 particles that show rod-like inner densities. Scale bar: 50 nm.

**Fig. S8.**
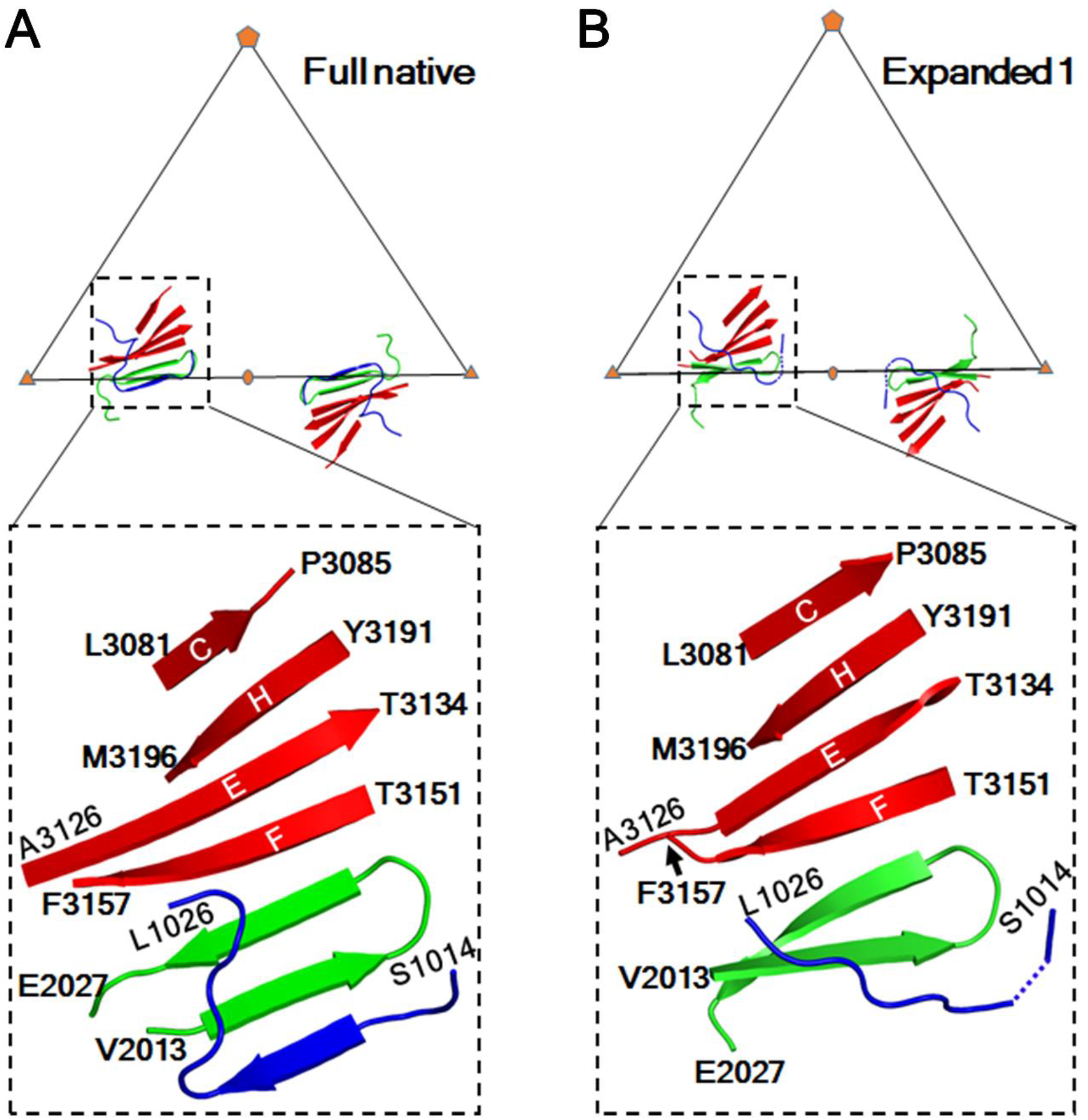
The seven-stranded interpentamer sheets in full native virions ***(A)*** are partially disrupted in E1 particles ***(B)***. Each triangle indicates an icosahedral asymmetric unit. VP1, VP2, and VP3 are colored blue, green, and red, respectively. In each panel, a rectangle (dashed line) in the upper triangle defines the limit of the close-up view in the large bottom rectangle (dashed line), which is slightly tilted for better visualization. The β-strands C, H, E, and F in VP3 are labelled by their corresponding letters. Residues 1016-1019 are disordered in the E1 particle structure.

**Fig. S9.**
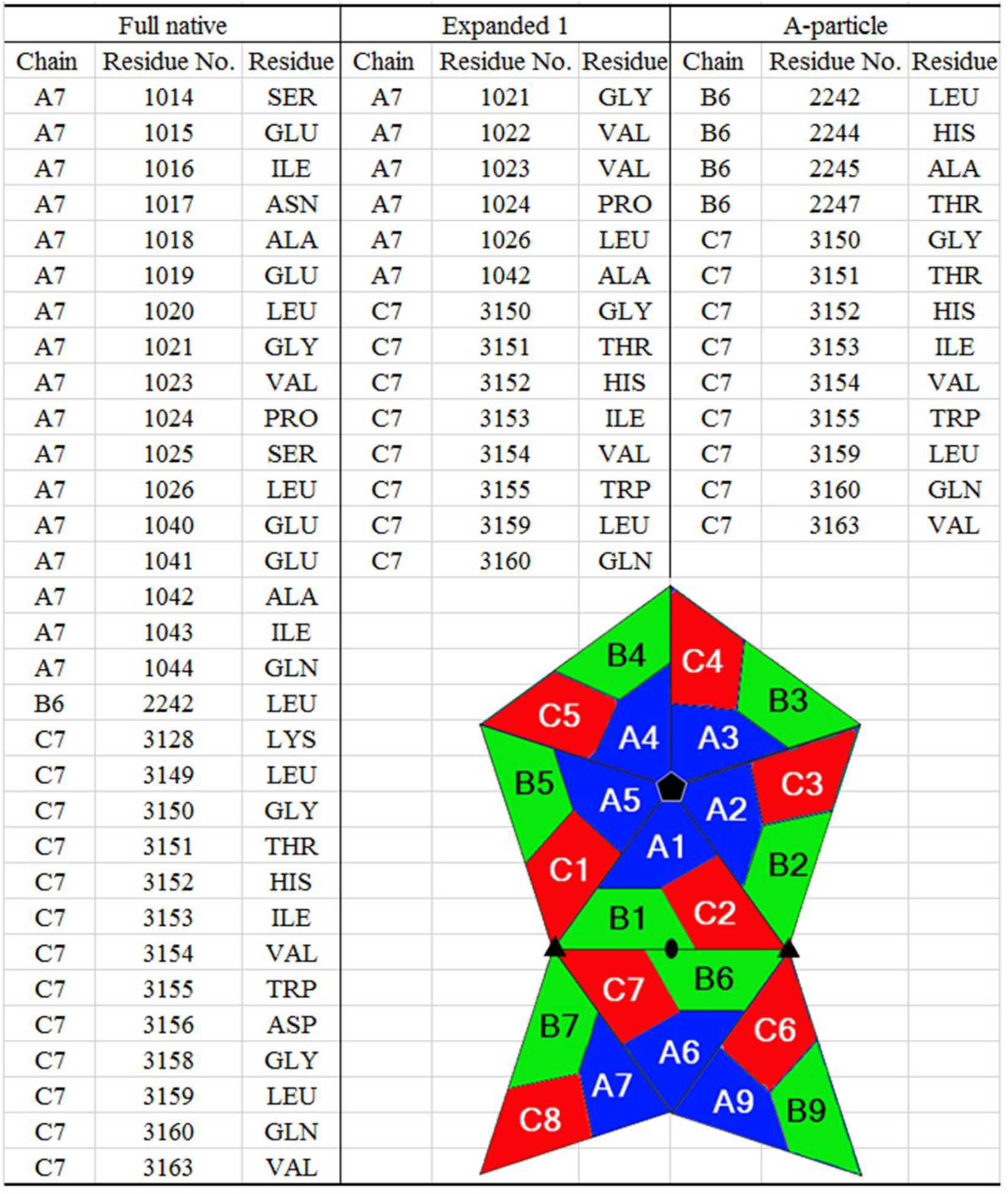
Structural reorganization of the VP2 N-termini weakens the interpentamer contact during uncoating. A schematic (bottom right) of eight icosahedral asymmetric units of the capsid is colored blue (VP1), green (VP2), and red (VP3). Each subunit is labelled by a chain number (e.g., A1). For each of the three full particle states (dataset B_4_Neu), the residues listed above are within a distance of 4 Å to any atom of residues 2013-2026 in chain B1.

**Fig. S10.**
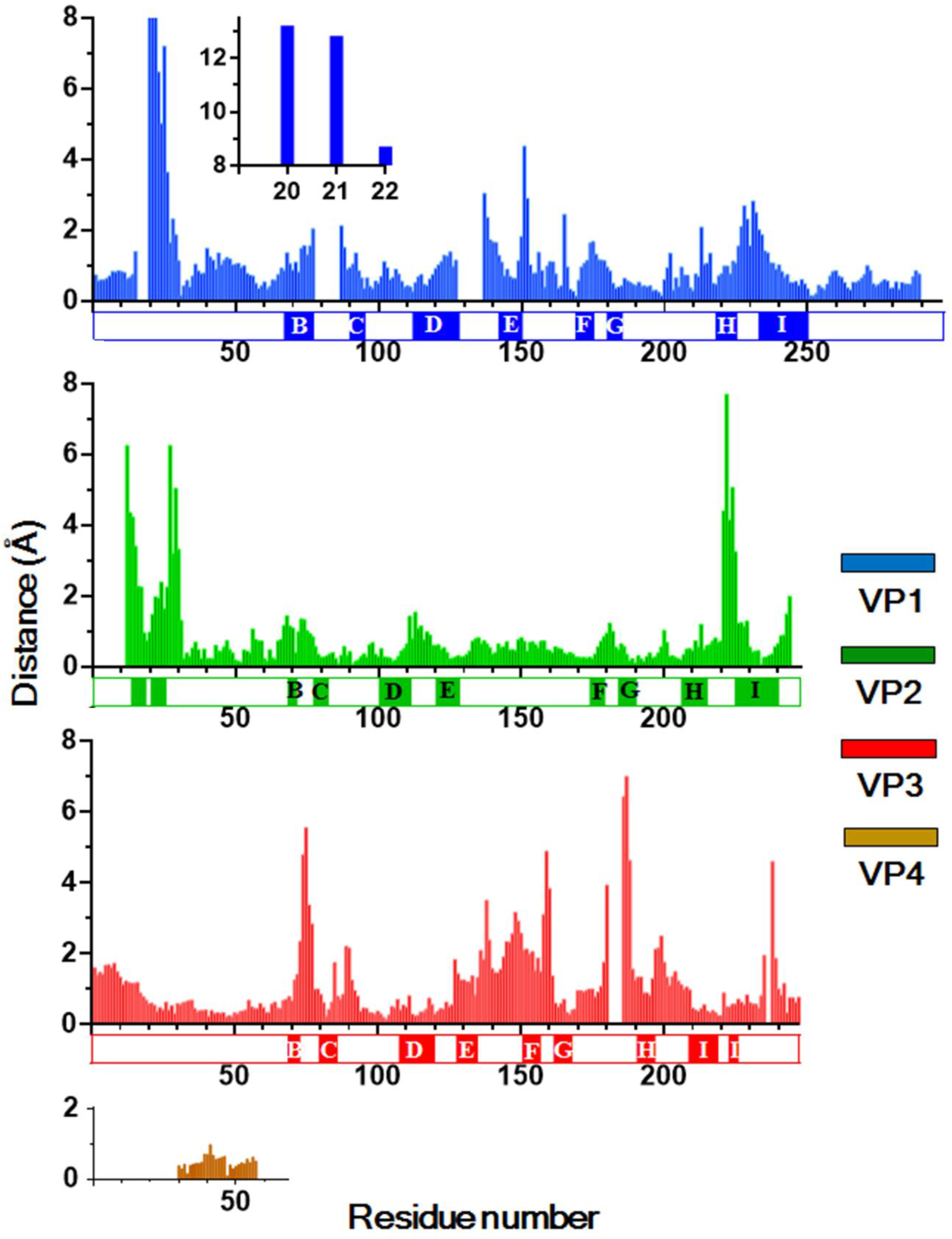
Distances between equivalent Cα atoms upon superposition of equivalent protomers in full native virions and E1 particles. Rectangular blocks denote residues that form β-strands. The jelly roll β-strands B, C, D, E, F, G, H, and I are labelled by their corresponding letters. The inset shows a close-up view of the plot for the VP1 N-terminal residues 1020-1022.

**Fig. S11.**
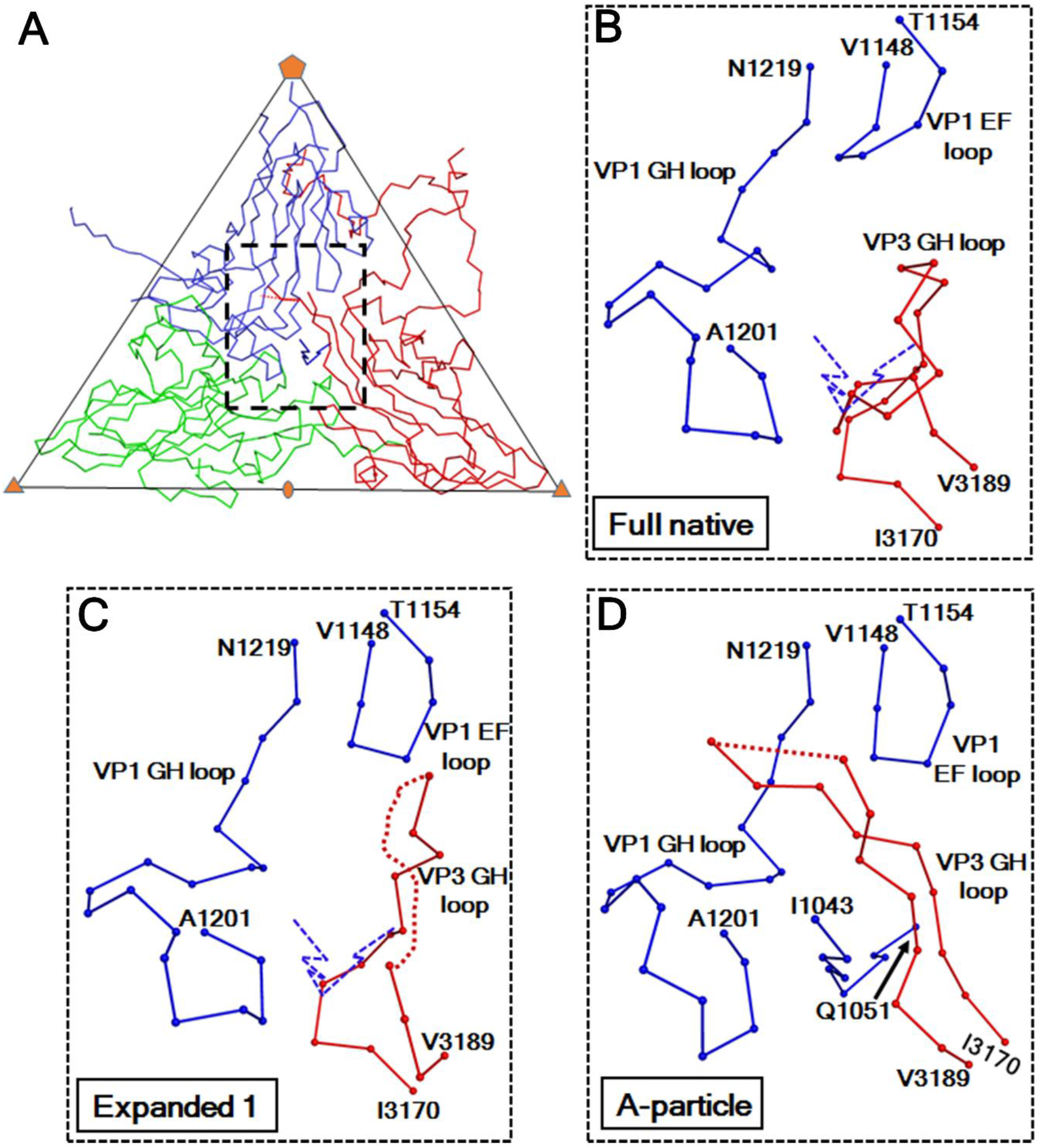
Structural rearrangements of loops near the quasi-three-fold axes facilitate the externalization of the VP1 N-termini. ***(A)*** The Cα backbone representation of an icosahedral asymmetric unit of the A-particle structure is colored blue (VP1), green (VP2), and red (VP3). The rectangle (dashed line) indicates the limit of the close-up views in ***(B-D)**. **(B)*** The structure of full native virions after superimposition of equivalent protomers in full native virions and A-particles. ***(C)*** The structure of E1 particles after superimposition of equivalent protomers in E1-particles and A-particles. In these two panels, a blue dashed line represents the Cα backbone trace of the VP1 N-terminal residues 1043-1051 observed in the A-particle structure. These residues would clash with the VP3 GH loop in full native virions and with that in E1 particles. The VP3 GH loop adopts a coiled conformation in full native virions. This loop becomes partially disordered and adopts a more extended conformation in the other two structural states.

**Fig. S12.**
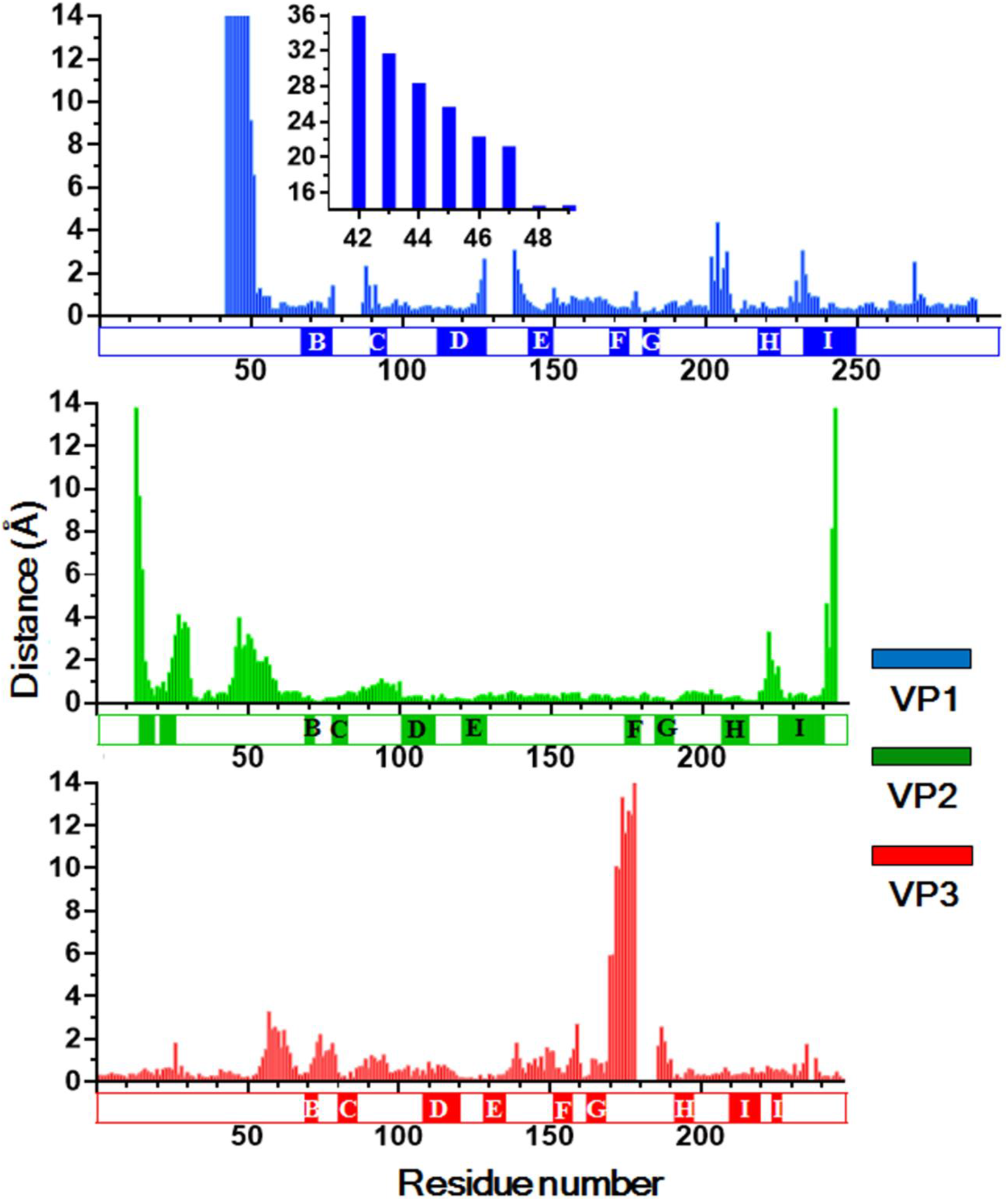
Distances between equivalent Cα atoms upon superposition of the equivalent protomers in E1 particles and A-particles. The rectangular blocks are defined as in **Fig. S11**. The inset shows a close-up view of the plot for the VP1 N-terminal residues 1042-1049.

**Table S1.** Cryo-EM data statistics.

|  | A_Native<br>(full) | B_RT_Acid<br>(A-particle) | B_RT_Acid<br>(emptied) | B_33_Acid<br>(A-particle) | B_33_Acid<br>(emptied) |
| --- | --- | --- | --- | --- | --- |
| **Data collection and processing<sup>a</sup>** |  |  |  |  |  |
| No. of micrographs<sup>b</sup> | 642 | 144 | 144 | 950 | 950 |
| Pixel size<sup>c</sup> (Å/pixel) | 0.65 | 0.65 | 0.65 | 0.65 | 0.65 |
| Dose rate (e-/pixel/s) | 8 | 8 | 8 | 8 | 8 |
| Total dose (e-/Å<sup>2</sup>) | 36 | 28 | 28 | 38 | 38 |
| Frame rate (ms) | 200 | 200 | 200 | 200 | 200 |
| Defocus (µm) | 0.3-3.0 | 0.9-4.8 | 0.9-4.8 | 0.5-3.5 | 0.5-3.5 |
| No. of ptcls for reconstruction | 16021 | 3708 | 2150 | 23082 | 19325 |
| Resolution<sup>d</sup> (Å) | 2.17 | 3.25 | 3.75 | 2.73 | 2.90 |
| Map sharpening B-factor (Å<sup>2</sup>) | -83.0 | -125.7 | -163.7 | -120.2 | -144.1 |
| **Model Statistics** |  |  |  |  |  |
| Correlation coefficient<sup>e</sup> | 0.838 | 0.834 | 0.829 | 0.848 | 0.846 |
| <u>No. of atoms</u> |  |  |  |  |  |
| Protein | 6822 | 5362 | 5025 | 5579 | 5083 |
| Water | 500 | 0 | 0 | 253 | 0 |
| Avg. B-factor (Å<sup>2</sup>) | 18.2 | 39.9 | 49.2 | 30.4 | 32.5 |
| <u>r.m.s deviations<sup>f</sup></u> |  |  |  |  |  |
| Bond lengths (Å) | 0.010 | 0.009 | 0.008 | 0.010 | 0.011 |
| Bond angles (°) | 1.355 | 1.296 | 1.162 | 1.327 | 1.261 |
| <u>Ramachadran plot<sup>f</sup></u> |  |  |  |  |  |
| Favored (%) | 96.1 | 93.9 | 91.5 | 94.7 | 93.6 |
| Allowed (%) | 3.6 | 5.9 | 8.2 | 5.0 | 5.9 |
| Outliers (%) | 0.3 | 0.2 | 0.3 | 0.3 | 0.5 |

<sup>a</sup> Cryo-EM data were collected using an FEI Titan Krios transmission electron microscope operated at 300 kV and equipped with a Gatan K2 Summit direct electron detector.
<sup>b</sup> Micrographs from which particles were selected.
<sup>c</sup> Super resolution pixel size. The physical pixel size is 1.3 Å/pixel
<sup>d</sup> Estimated based on Fourier shell correlation between two half maps using a cutoff of 0.143 (71)
<sup>e</sup> Between the cryo-EM map and a map calculated based on the atomic model specifying the cryo-EM map resolution
<sup>f</sup> Based on the criteria of MolProbity (78)

**Table S2.** Root-mean-square deviations (Å) between pairs of EV-D68 structures^a^.

|  | Native_full<sup>b</sup> | RT_A<sup>c</sup> | RT_EMP<sup>c</sup> | 33_A<sup>d</sup> | 33_EMP<sup>d</sup> |
| --- | --- | --- | --- | --- | --- |
| Native_full<sup>b</sup> | - |  |  |  |  |
| RT_A<sup>c</sup> | 5.7 | - |  |  |  |
| RT_EMP<sup>c</sup> | 4.6 | 0.8 | - |  |  |
| 33_A<sup>d</sup> | 5.7 | 0.6 | 0.7 | - |  |
| 33_EMP<sup>d</sup> | 4.6 | 0.7 | 0.6 | 0.5 | - |

<sup>a</sup> Calculated based on equivalent Cα atoms when icosahedral symmetry axes are aligned.
<sup>b</sup> Dataset A\_Native
<sup>c</sup> Dataset B\_RT\_Acid. A and EMP refer to A-particle and emptied particle, respectively.
<sup>d</sup> Dataset B\_33\_Acid. A and EMP refer to A-particle and emptied particle, respectively.

**Table S3.**
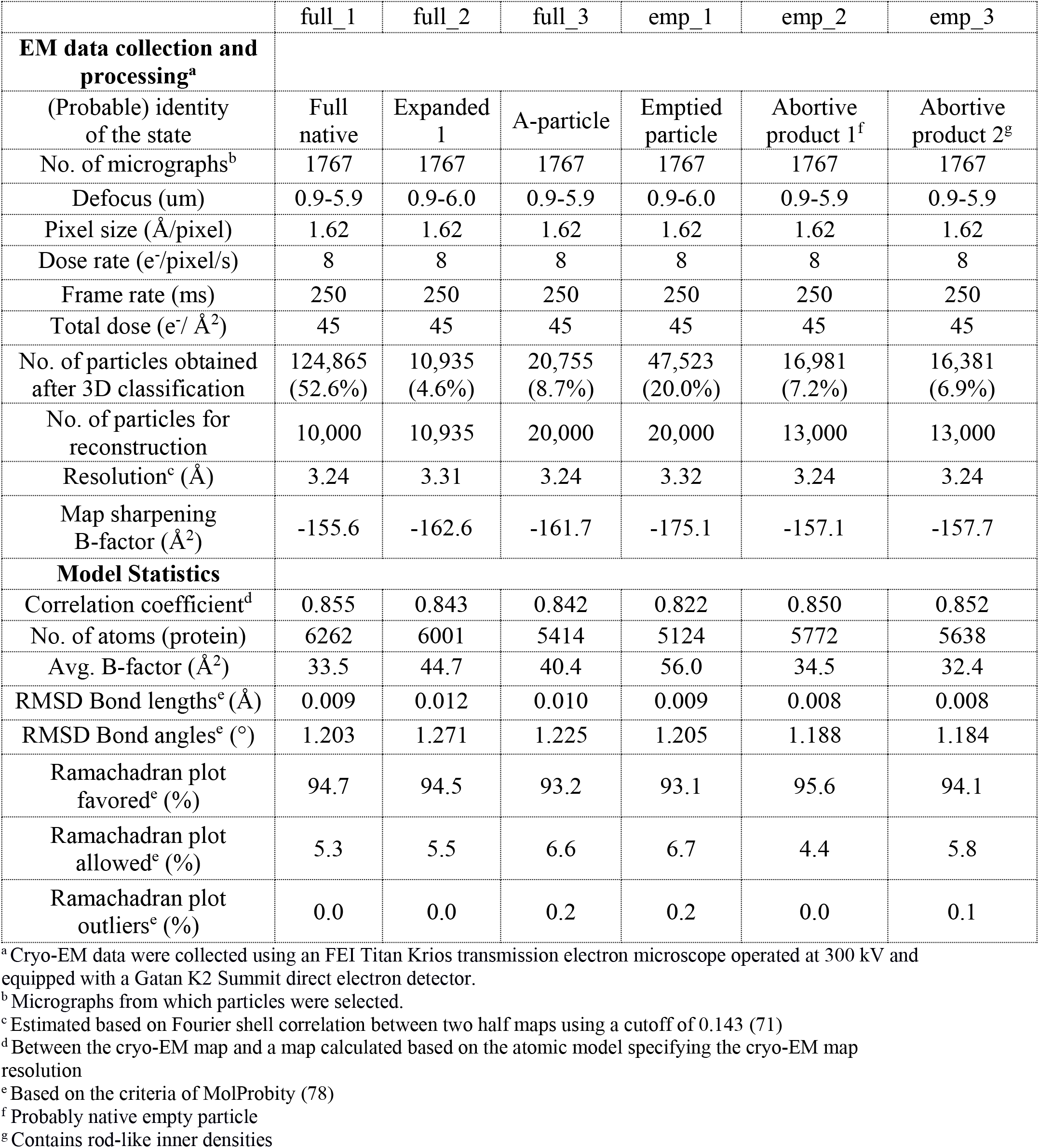
Cryo-EM data statistics for dataset B_4_Neu.

**Table S4.** Root-mean-square deviations (Å) between pairs of structures (dataset B_4_Neu).^a^.

|  | Full<br>native | Expanded 1 | A-particle | Emptied<br>particle | Abortive<br>product 2 | Abortive<br>product 1 |
| --- | --- | --- | --- | --- | --- | --- |
| Full native | - |  |  |  |  |  |
| Expanded 1 | 4.4 | - |  |  |  |  |
| A-particle | 6.0 | 3.3 | - |  |  |  |
| Emptied particle | 4.9 | 1.1 | 1.1 | - |  |  |
| Abortive product 2 | 1.0 | 3.5 | 4.3 | 4.0 | - |  |
| Abortive product 1 | 1.1 | 3.5 | 4.3 | 4.0 | 0.3 | - |

<sup>a</sup>Calculated based on equivalent Cα atoms when icosahedral symmetry axes are aligned.

**Table S5.** Rigid body movements of capsid proteins when full native virions are converted into E1 particles.

| Protein | Psi<sup>a</sup> (°) | Phi<sup>a</sup> (°) | Kappa<sup>b</sup> (°) | Translation (Å) | Rmsd<sup>c</sup> (Å) |
| --- | --- | --- | --- | --- | --- |
| VP1 | 77.0 | 38.0 | 2.8 | 4.5 | 1.1 |
| VP2 | 83.5 | 17.5 | 3.3 | 3.4 | 1.3 |
| VP3 | 71.6 | 15.2 | 2.8 | 3.9 | 1.4 |
| VP4 | 48.7 | 40.8 | 3.3 | 4.7 | 0.3 |

<sup>a</sup> These two polar angles define the direction of the rotational axis according to (82)
<sup>b</sup> Clockwise rotation as viewed out from inside the virus.
<sup>c</sup> Between equivalent Cα atoms

**Table S6.**
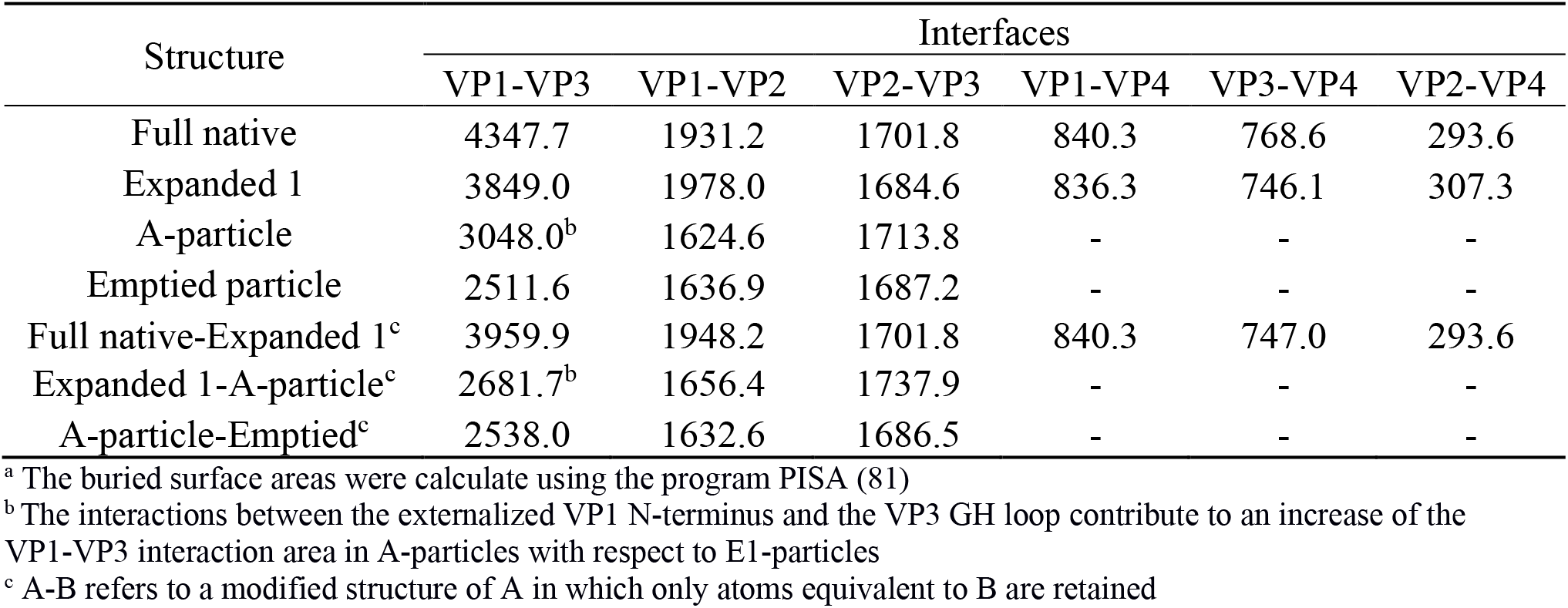
Buried surface areas (Å^2^) at the intra-protomer protein-protein interacting interfaces.^a^.

**Table S7.**
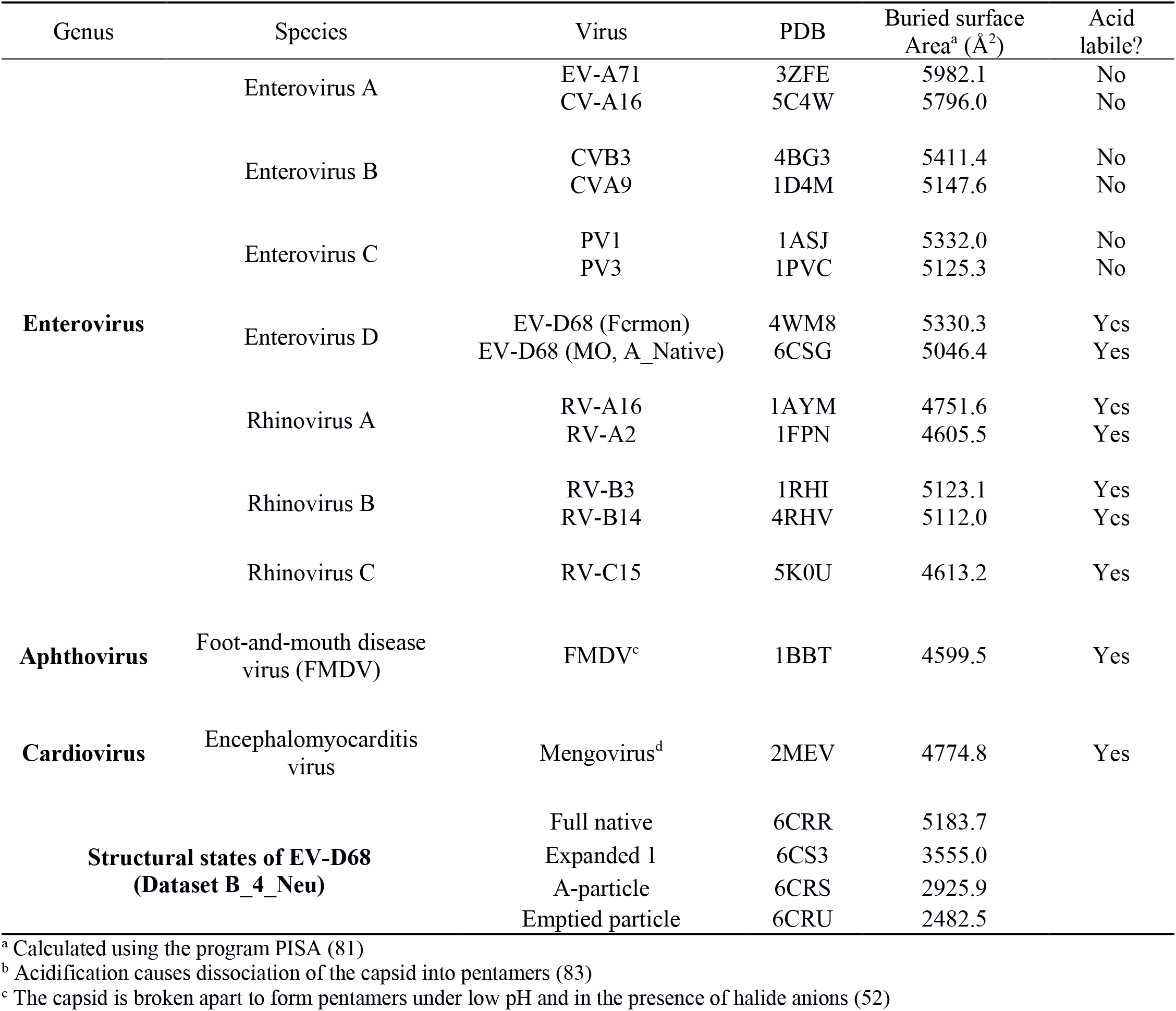
Comparison of buried surface areas at the interpentamer interfaces.

| Genus | Species | Virus | PDB | Buried surface Area<sup>a</sup> (Å<sup>2</sup>) | Acid labile? |
| --- | --- | --- | --- | --- | --- |
| **Enterovirus** | Enterovirus A | EV-A71 | 3ZFE | 5982.1 | No |
|  |  | CV-A16 | 5C4W | 5796.0 | No |
|  | Enterovirus B | CVB3 | 4BG3 | 5411.4 | No |
|  |  | CVA9 | 1D4M | 5147.6 | No |
|  | Enterovirus C | PV1 | 1ASJ | 5332.0 | No |
|  |  | PV3 | 1PVC | 5125.3 | No |
|  | Enterovirus D | EV-D68 (Fermon) | 4WM8 | 5330.3 | Yes |
|  |  | EV-D68 (MO, A_Native) | 6CSG | 5046.4 | Yes |
|  | Rhinovirus A | RV-A16 | 1AYM | 4751.6 | Yes |
|  |  | RV-A2 | 1FPN | 4605.5 | Yes |
|  | Rhinovirus B | RV-B3 | 1RHI | 5123.1 | Yes |
|  |  | RV-B14 | 4RHV | 5112.0 | Yes |
|  | Rhinovirus C | RV-C15 | 5K0U | 4613.2 | Yes |
| **Aphthovirus** | Foot-and-mouth disease virus (FMDV) | FMDV<sup>c</sup> | 1BBT | 4599.5 | Yes |
| **Cardiovirus** | Encephalomyocarditis virus | Mengovirus<sup>d</sup> | 2MEV | 4774.8 | Yes |
| **Structural states of EV-D68 (Dataset B_4_Neu)** |  | Full native | 6CRR | 5183.7 |  |
|  |  | Expanded 1 | 6CS3 | 3555.0 |  |
|  |  | A-particle | 6CRS | 2925.9 |  |
|  |  | Emptied particle | 6CRU | 2482.5 |  |

<sup>a</sup> Calculated using the program PISA (81)
<sup>b</sup> Acidification causes dissociation of the capsid into pentamers (83)
<sup>c</sup> The capsid is broken apart to form pentamers under low pH and in the presence of halide anions (52)

**Table S8.** Differences of amino acid sequences in capsid proteins between EV-D68 strains used in this work.

| Residue No.<sup>a</sup> | Strain Fermon | Strains MO/KY |
| --- | --- | --- |
| 1072 | LYS | ARG |
| 1085 | GLY | ARG |
| 1087 | HIS | ASP |
| 1132 | ASP | ASN |
| 1156 | LYS | GLU |
| 1157 | GLU | LYS |
| 1270 | ARG | LYS |
| 1271 | ASP | GLU |
| 1272 | THR | ARG |
| 1288 | THR | LYS |
| 2067 | ARG | LYS |
| 2098 | TYR | HIS |
| 2136 | ASP | ASN |
| 2144 | ASN | ASP |
| 2151 | ARG | GLU |
| 3059 | ASP | GLU |
| 3068 | ARG | LYS |

<sup>a</sup> Numbering based on the amino acid sequence of EV-D68 strain Fermon

**Table S9.** Spatial and temporal origins of EV-D68 strains that have a tyrosine at position 2098.

| GenBank accession No. | Country | Year of collection |
| --- | --- | --- |
| AMQ48961.1 | USA | 1962 |
| AMQ48960.1 | USA | 1962 |
| AAR98503.1 | USA | 1962 |
| ALG02131.1 | USA | 1963 |
| AKQ43512.1 | Netherlands | 2009 |
| ANJ61749.1 | USA | 2009 |
| ANJ61729.1 | USA | 2009 |
| ANJ61595.1 | USA | 2009 |
| ANJ61561.1 | USA | 2009 |
| ANJ61547.1 | USA | 2009 |
| ANJ61542.1 | USA | 2009 |
| AFM73548.1 | USA | 2009 |
| AKQ43520.1 | Netherlands | 2010 |
| AKQ43518.1 | Netherlands | 2010 |
| AGC00381.1 | New Zealand | 2010 |
| AOR17470.1 | Philippines | 2011 |
| ANJ61563.1 | USA | 2011 |
| ANJ61590.1 | USA | 2012 |
| ANJ61582.1 | USA | 2012 |
| ANJ61578.1 | USA | 2012 |
| ANJ61568.1 | USA | 2012 |
| ANJ61567.1 | USA | 2012 |
| ANJ61556.1 | USA | 2012 |
| ANJ61548.1 | USA | 2012 |
| ANJ61546.1 | USA | 2012 |
| ANJ61545.1 | USA | 2012 |
| AKU75647.1 | USA | 2012 |
| BAW35377.1 | Japan | 2013 |
| BAW35376.1 | Japan | 2013 |
| BAW35375.1 | Japan | 2013 |
| BAW35374.1 | Japan | 2013 |

